# LRRK2 and Rab10 Coordinate Macropinocytosis to Mediate Immunological Responses in Phagocytes

**DOI:** 10.1101/2020.01.30.926840

**Authors:** Zhiyong Liu, Enquan Xu, Hien Tran Zhao, Tracy Cole, Andrew B. West

**Author notes:** C.A. Information, 3 Genome Court, Durham, North Carolina, 27710 USA.

## Abstract

Genetic variation in *LRRK2* associates with susceptibility to Parkinson’s disease, Crohn’s disease, and mycobacteria infection, with high expression of LRRK2, and the LRRK2 kinase substrate Rab10, in phagocytic cells in the immune system. In mouse and human primary monocyte-derived macrophages, dendritic cells, and microglia-like cells, we find that Rab10 specifically regulates a specialized form of endocytosis known as macropinocytosis, without affecting phagocytosis or clathrin-mediated endocytosis. LRRK2 phosphorylates cytoplasmic PI(3,4,5)P_3_-positive GTP-Rab10 early macropinosomes, before EEA1 and Rab5 recruitment occurs. Macropinosome cargo in macrophages includes CCR5, CD11b, and MHCII, with LRRK2-phosphorylation of Rab10 potently blocking EHBP1L1-mediated recycling tubules and cargo turnover. EHBP1L1 over-expression competitively inhibits LRRK2-phosphorylation of Rab10, mimicking the effects of LRRK2 kinase inhibition in promoting cargo recycling. Both Rab10 knockdown and LRRK2 kinase inhibition potently suppresses the maturation of macropinosome-derived CCR5-loaded signaling endosomes important for CCL5-induced AKT-activation and chemotaxis. These data support a novel axis in the endolysosomal system whereby LRRK2-mediated Rab10 phosphorylation stalls vesicle fast-recycling to promote PI3K-AKT signal transduction.

## INTRODUCTION

Macropinocytosis, a process originally discovered in macrophages isolated from rats ^1^, is a clathrin-independent form of endocytosis with many similarities to the better understood process of phagocytosis ^2^. Both phagocytosis and macropinocytosis are actin-dependent, with phagocytosis mediating internalization of large and insoluble cargo and macropinocytosis supporting internalization of small and soluble particles. Constitutive macropinocytosis occurs in monocytes, macrophages, and other phagocytic cells, and may be critical for immune surveillance ^2^, although specific signaling pathways regulated by macropinocytosis are not well understood. Macropinocytosis can occur in non-phagocytic cells as well, usually in acute response to growth factors. Possible roles include the uptake and turn-over of misfolded proteins in Parkinson’s disease (PD), Alzheimer’s disease (AD), and amyotrophic lateral sclerosis (ALS) ^3–5^. Without knowledge of proteins that specifically regulate macropinocytosis in endocytosis, most studies have explored the process using small molecule inhibitors EIPA (5-(N-Ethyl-N-isopropyl)amiloride) and rottlerin ^6, 7^. However, while both molecules efficiently block macropinocytosis, they also affect other forms of endocytosis ^2^.

Rab small GTPases can selectively control endocytic processes, vesicle formation, trafficking and maturation, and recycling and degradation ^8, 9^. In phagocytosis, Rab35 drives phagosome formation, whereas Rab20 is critical in phagosome maturation ^10^. Rab proteins specific for macropinocytosis, macropinosome formation, and macropinosome maturation, have not previously been identified. The *Rab10* gene is linked to human disease in several ways. Genetic variants in *Rab10* associate with resilience to AD, and a phosphorylation site in the Rab10 switch II domain is upregulated by PD-associated mutations in the LRRK2 protein kinase ^11^. LRRK2-mediated phosphorylation of Rab10 is strongly affected by all known pathogenic mutations as well as PD-risk factor genetic variants like G2385R, with Rab10 phosphorylation changing the affinity of different binding proteins to the Rab10 effector loops ^12^. Our past work, suggests LRRK2 phosphorylation of Rab10 may prolong Rab10 in a GTP-bound state by inhibiting Rab10 interactions with GAP proteins ^13^. While the specific role of Rab10 in cells linked to neurodegeneration is under investigation, recent work demonstrates Rab10 downregulates ciliogenesis in some types of cells and neurons in the brain ^14^. Further, in COS-7 cells, Rab10 may regulate ER dynamics and morphology ^15^. Finally, Rab10 may facilitate the traffic of toll-like receptor (TLR) complexes from the Golgi to the plasma membrane in macrophages exposed to lipopolysaccharides (LPS) ^16^. This regulation may have relevance for TLRs important in binding protein fibrils associated with neurodegenerative diseases ^17^.

Herein, we find Rab10 expression drives the formation and maturation of macropinosomes in primary human and mouse phagocytic cells, without affecting other types of endocytosis or TLR4 signaling. While Rab10-positive macropinosomes normally rapidly recycle, locking Rab10 in a GTP-bound state stalls the vesicles in the cytoplasm that forces vesicle maturation to LAMP1-positive endosomes. LRRK2-mediated phosphorylation of Rab10 mimics GTP-locked Rab10 through blocking the actions of EHBP1L1-mediated recycling. Through immunostaining approaches, we found that nearly all macrophage macropinosomes are transport vesicles for the GPCR-chemokine receptor C-C chemokine receptor type 5 (CCR5). Reducing Rab10 expression, or blocking LRRK2-mediated Rab10 phosphorylation, inhibits CCR5 dependent AKT activation and chemotaxis. Together, these results provide evidence that Rab10 critically and specifically drives macropinocytosis in phagocytes. Manipulation of Rab10 and LRRK2 provides a new avenue to better understand constitutive vesicle internalization via macropinocytosis in controlling different immunological responses.

## RESULTS

### Rab10 knockdown impairs macropinocytosis without affecting phagocytosis or clathrin-mediated endocytosis

Recent RNA sequencing and single-cell sequencing approaches have highlighted selective expression of Rab10 in phagocytes, particularly in monocytes and macrophages (Supplemental Figure 1). The only known protein kinase to phosphorylate Rab10, LRRK2, shares a similar restricted expression profile in immune cell subtypes (Supplemental Figure 1). In our recent contributions to the development of validated monoclonal antibodies directed to Rab10 (validated with Rab10 knockdown, Supplemental Figure 2), we localized Rab10 to well defined patches of protein in macrophage plasma membrane ruffles. These ruffles persisted into clear (but short-lived) detached Rab10-positive vesicles in the cytoplasm (Supplemental Figure 2). This process in macrophages, highlighted by Rab10, appeared structurally reminiscent of constitutive macropinocytosis, a poorly understood endocytosis pathway inherent to phagocytic cells.

To determine if Rab10 knockdown affects macropinocytosis in phagocytic cells, we first optimized effective knockdown strategies in macrophages that did not activate or induce morphological changes in the cells. Incubation of cells with low micromolar concentrations of 3^rd^ generation stabilized anti-sense oligonucleotides (ASOs) efficiently reduced Rab10 expression in primary cells without macrophage activation (Table 1). Rab10-directed ASOs reduced expression to ∼10% of constitutive levels after two-days post ASO-exposure without affecting the expression of proteins from the same Rab-type I family including Rab8a, Rab7, Rab5, or Rab3a (Figure 1A,B). Feeding macrophages 70 kDa fluorescent dextran, a canonical marker used to highlight macropinosomes, or IgG-beads (used to highlight phagosomes), or fluorescent transferrin, revealed a severe impairment only in dextran uptake due to Rab10 deficiency (Figure 1C-F). Other morphological abnormalities in the plasmam membrane or endolysosomal system due to Rab10 deficiency were not observed. We applied the same Rab10 knockdown strategy to both human and mouse primary monocyte-derived cells polarized to macrophages (treated with M-CSF), dendritic cells (treated with GM-CSF), or microglia-like cells (treated with IL-34, NGF-β, CCL2), and in each case dextran-uptake was specifically inhibited, without apparent effect in other cell compartments. The degree of reduction of macropinocytosis caused by Rab10 deficiency was comparable to micromolar-exposures of the small molecule macropinocytosis inhibitor EIPA (Figure 1G-N). Results and effect sizes with Rab10 knockdown in impairing macropinocytosis were robustly replicated in several other phagocyte cell lines and primary-cultured cells from both human and mouse, and with a second independent-targeting ASO (Supplemental Figure 3).

**Figure 1.**
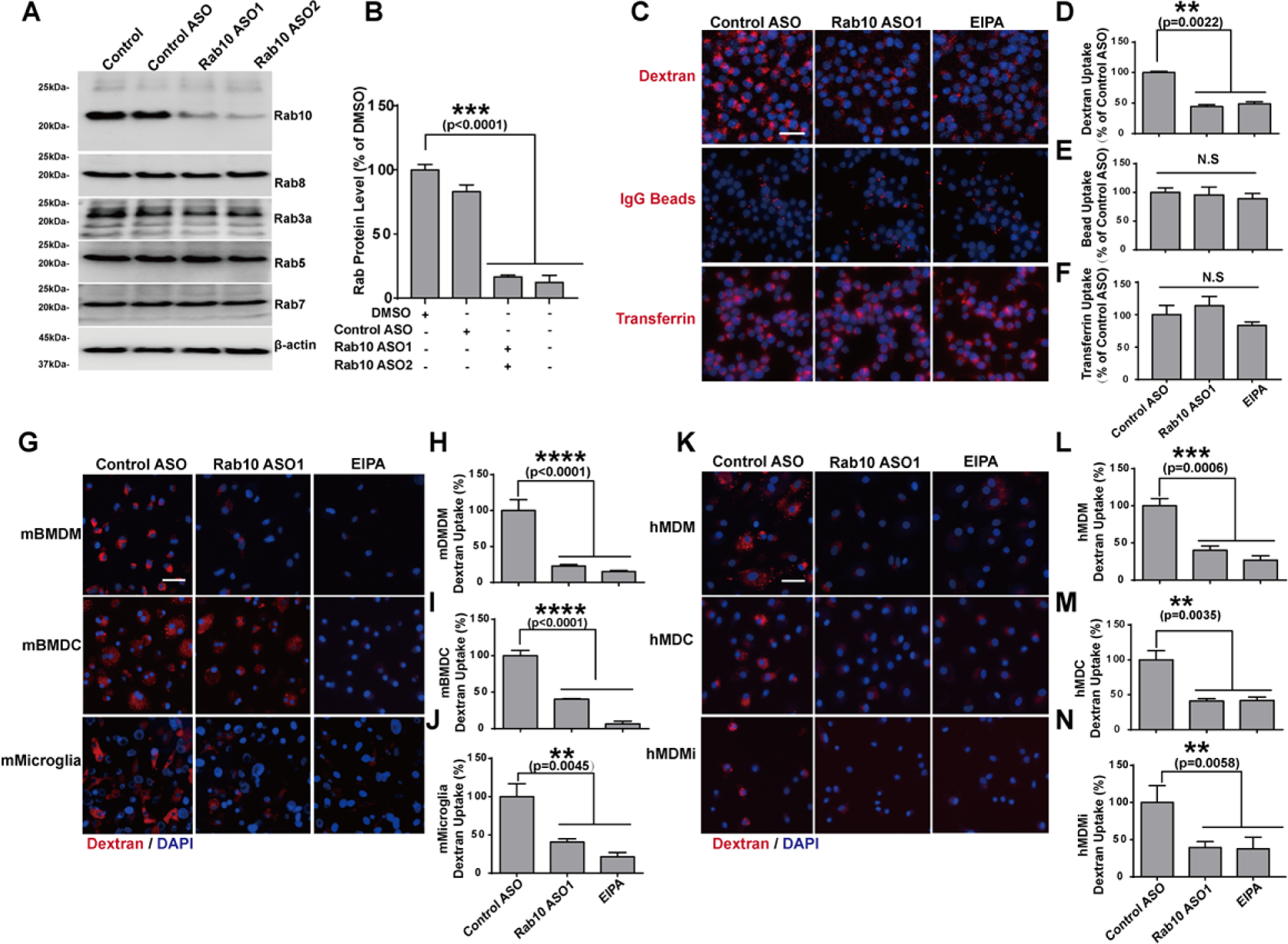
Rab10 expression regulates macropinocytosis. **A)** Raw 264.7 macrophage cells were treated with DMSO (0.05%), 1 μM control antisense oligonucleotide (ASO), or Rab10-directed ASO (1 μM), for four days. Immunoblots are representative of three independent experiments, with similar results obtained. **B)** Calculated reduction of Rab10 (n=3 biologically independent experiments). **C)** Raw 264.7 cells were treated with the indicated ASO for four days, or 50 μM EIPA (5-[N-ethyl-N-isopropyl]amiloride) for 20 min, prior to incubation for 60 min with fluorescent (tetramethylrhodamine, TRITC)-labeled dextran (70 kDa, shown as red color), IgG-agarose, or transferrin. Representative photomicrographs (from >30 images analyzed for each conditions from n=3 biologically independent experiments) are shown from cells extensively washed, fixed and stained with DAPI (shown as blue color). **D-F)** Relative fluorescent signals (i.e., dextran uptake) were calculated as a percent of control-ASO treated cells. **G)** Primary mouse bone marrow-derived macrophage cells (BMDM) from adult male C57BL/6J mice were treated with murine cytokines directed towards a monocytic lineage (mBMDM), dendritic lineage (mBMDC), or microglia-like lineage (mMicroglia). Cells were incubated with TRITC-dextran for 30 min prior to washing, fixing, and staining with DAPI. Representative photomicrographs from >30 images analyzed for each condition from n=3 biologically independent experiments are shown. **H-J),** Relative fluorescent signals were calculated as a percent of control-ASO treated cells. **K)** Human monocytes purified from venous blood draws from healthy adult males were treated with human cytokines towards a monocytic lineage (hMDM), dendritic lineage (hMDC), or microglia-like lineage (hMicroglia). Cells were incubated with TRITC-dextran for 30 min prior to washing, fixing, and staining with DAPI. Representative photomicrographs from >30 images analyzed for each conditions from n=3 biologically independent experiments are shown. **L-N)** Relative fluorescent signals were calculated as a percent of control-ASO treated cells. Scale bars show 50 μm. Column graphs show group means with error bars as ± SEM; significance was assessed by one-way ANOVA (p>0.01) with * representing Tukey’s post hoc test p<0.01, or n.s. (p>0.05).

**Table 1.** Sequences of antisense oligos. Black: unmodified deoxyribose (2’H); orange: 2’ methoxyethyl (MOE). Unmarked linkages: phosphorothioate (PS); linkages marked with o: normal phosphodiester (PO). mC: 5-methylcytosine.

| **ASO** | **Sequence** |
| --- | --- |
| Control ASO "Ionis 676630" (CTRL) | mCmCoToAoTAGGAmCTATmCmCAoGoGoAA |
| Rab10 "Ionis 978453" (ASO1) | TmComCGoAAATATGTGGTAoGoTAmC |
| Rab10 "Ionis 978554" (ASO2) | GToTToCAGGATATGATmCoGoGmCT |

### Rab10 regulates the early trafficking of macropinosome

To pinpoint Rab10 initial entry into ruffled membranes, co-localization experiments with the PLCδ-PH-domain revealed initial exclusion of Rab10 from the beginning of circular cupping known to be regulated by ARF and Rac1-GTPases (Figure 2A,B) ^18^. Only sparse Rab10 protein could be localized to dextran-loaded SARA-FYVE domain-positive vesicles, known to be regulated in part by Rab21 and Rab5-GTPases in endocytosis ^19^. In contrast, nearly all Rab10-positive dextran-loaded vesicles were labeled completely labeled with the AKT-PH domain, suggesting Rab10 is specifically and rapidly recruited to the PIP_3_/PI(3,4,)P_2_ lipids on macropinosomes. Accordingly, brief application of wortmannin, a phosphatidylinositol 3-kinase inhibitor, completely ablated the ability of Rab10 to interact with vesicles (Figure 2C). In transfected Raw264.7 cells, live-cell imaging experiments revealed lifetimes of total Rab10 vesicles of less than ten minutes, with a subset of macropinosomes dropping most eGFP-Rab10 expression in exchange for mRuby-Rab5 in about eight minutes (Figure 2D, E). These results show that Rab10 is recruited to early (but not initial) stages of macropinocytosis, before Rab5 (an early endosome marker) is recruited.

**Figure 2.**
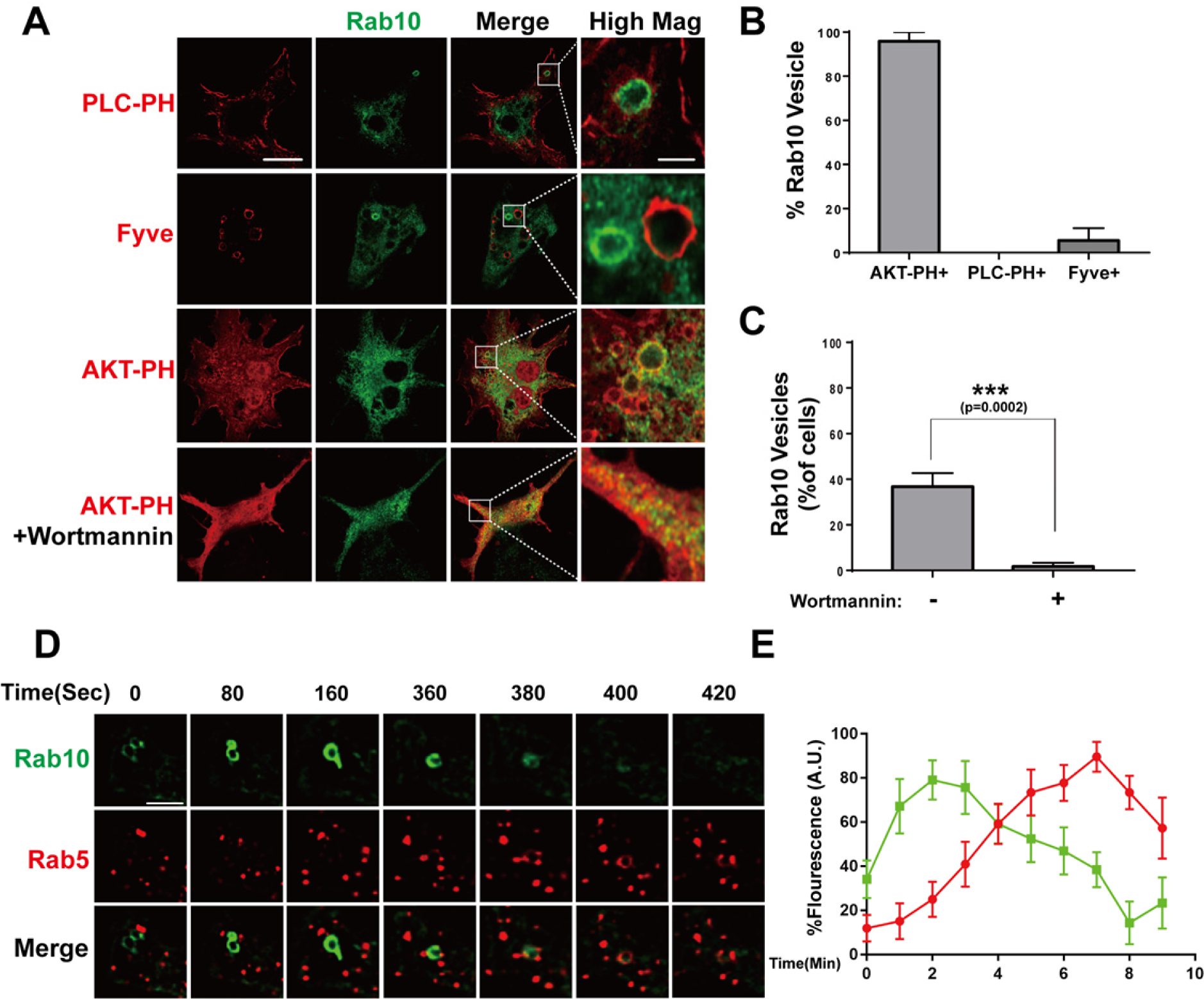
Rab10 is recruited to early plasma-membrane PIP_3_/ PI(3,4)P_2_ macropinsomes. **A)** Raw 264.7 macrophage cells were transfected with FLAG(N-term)-Rab10 and eGFP(C-term)-AKT-PH domain (PI(3,4,5)P_3_ marker), or eGFP(C-term)-SARA-Fyve domain (PI3P marker), or eGFP(C-term)-PLC-PH domain (PI(4,5)P_2_ marker), and treated with or without wortmannin (1 uM) for one hour. Representative photomicrographs (from >30 images analyzed for each conditions from n=3 biologically independent experiments) are shown from fixed cells immuno-stained for Flag-tag (Rab10, shown as green) together with eGFP epifluorescence (shown as red). White bounding boxes show “High Mag” panels that magnify representative individual vesicles. Scale bars are 10 μm or 1 μm in “Hig Mag” panels. **B)** Calculated percentage of double-positive vesicles (n>30 counted from n=3 biologically independent experiments) are shown. **C)** Percent of cells in the same photomicrographs harboring Rab10 vesicles are calculated. Significance is assessed by Student’s t-test with *** represents p<0.001. **D)** Raw 264.7 were co-transfected with eGFP(N-term)-Rab10 and mRuby(N-term)-Rab5 for 48 hours, with live-cell recordings for ∼10 minutes. A representative vesicle is shown. Scale bar represents 1 μm. **E)** Calculated relative fluorescence over time is indicated from Rab10 positive vesicles, initially emergent from the plasma membrane, that become positive for Rab5. Data show n=8 recorded vesicles across n=3 biologically independent experiments. Error bars show ± SEM.

GTP-hydrolysis activity of Rab GTPases can control localization. To study how GTP hydrolysis activity affects the localization and function of Rab10, macrophage cells were transfected with WT or GTP-locked Rab10 (Q68L), or a GDP-locked version (T23N). Consistent with live cell imaging, WT-Rab10 predominantly localizes on dextran filled vesicles in the absence of EEA1 (another early endosome marker) known to be the effector of Rab5 (Figure 3). In contrast, macrophage cells transfected with Q68L-Rab10 showed increased numbers of enlarged Rab10 positive macropinosomes, all filled with dextran but now stalled in the cytoplasm. These stalled Q68L-Rab10 loaded macropinosomes were degradative vesicles, evidenced by BODIPY ovalbumin fluorescence with LAMP1 reactivity (Figure 3). In contrast, GDP-locked Rab10 failed to associate with any vesicles in the cytoplasm. These results show that GTP binding is required for the macropinosome localization of Rab10, while GTP hydrolysis activity is required for the dissociation of Rab10 from macropinosomes. Q68L-Rab10 localization specifically to dextran loaded vesicles provides a second link, independent from Rab10 deficiency, to Rab10 mediation of macropinocytosis.

**Figure 3.**
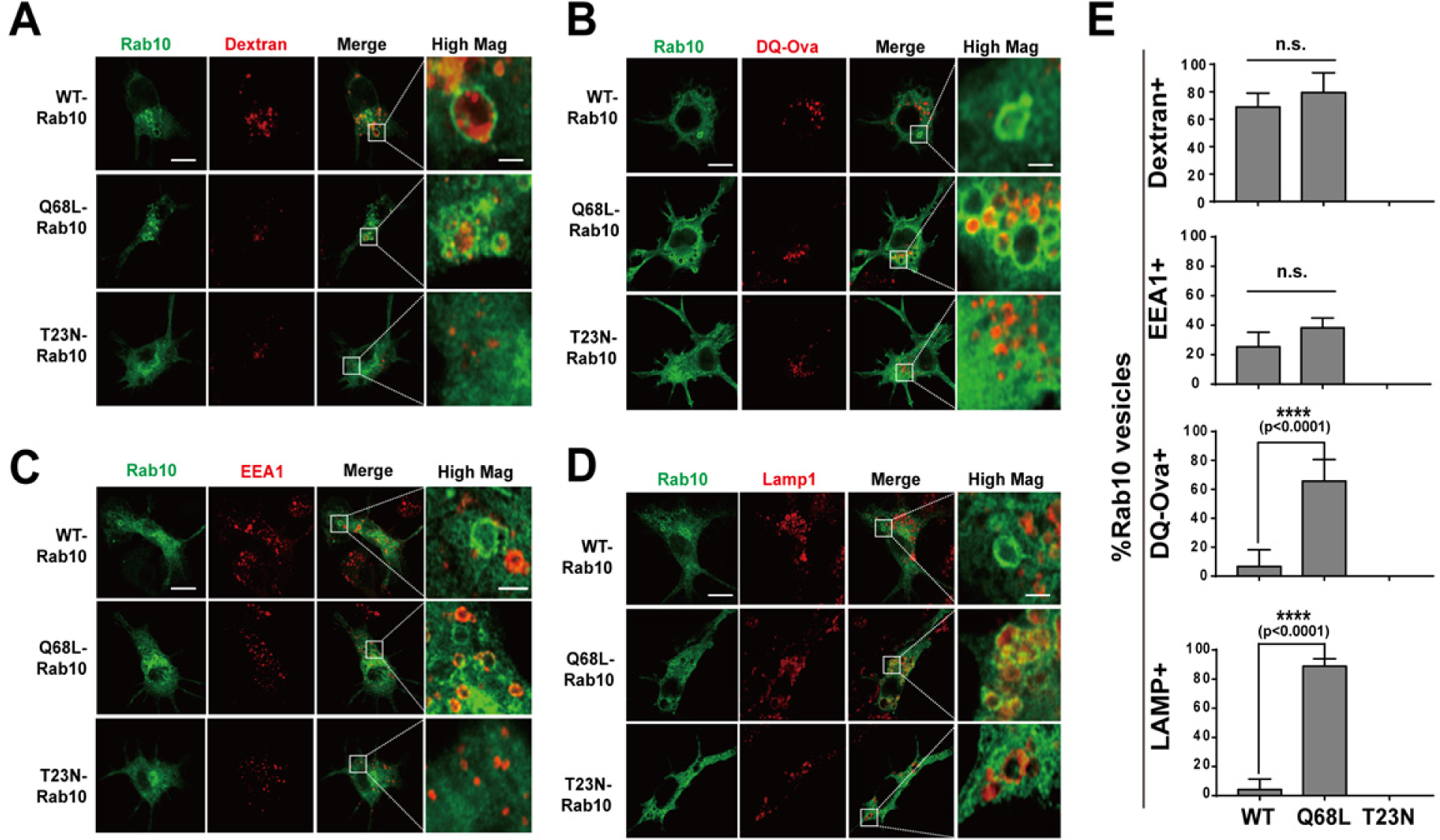
GTP-locked mutation stalls Rab10 on macropinosomes. **A)** Raw264.7 cells were transfected with plasmids expressing eGFP(N-term)-WT-Rab10, Q68L-Rab10 (GTP-locked) or T23N-Rab10 (GDP-locked). 24-hours later, cells were incubated with TRITC-dextran (70 kDa, shown as red signal), or **B)** DQ-ovalbumin, for 30 min before washing and fixation. **C)** Cells were further stained for EEA1, or, **D)** LAMP1, which was detected with Cy5 dye (show as red signal). Representative photomicrographs are from >30 cells analyzed for each condition from n=3 biologically independent experiments. White bounding boxes are magnified in “High Mag” panels that show individual vesicles. Scale bars represent 10 μm and 1 μm for “High Mag”. **E)** Calculated percentage of double-positive vesicles (n>30 vesicles counted from n=3 biologically independent experiments) are shown with error bars as ± SEM; significance was assessed by one-way ANOVA with **** representing Tukey’s post hoc test p<0.0001, or n.s. (p>0.05).

### LRRK2 phosphorylates Rab10 on early immature macropinosomes

Our previous experiments, consistent with others ^12–14^, reveal that a proportion of membrane associated GTP-bound Rab10 protein in transfected cell lines is phosphorylated by LRRK2 in the switch II loop at residue T73, exquisitely dependent on LRRK2 kinase activity in these cells. Consistent with our previous results, immunofluorescence staining using a pT73-Rab10 specific antibody confirmed that pT73-Rab10 signal exclusively localized to vesicular structures, as opposed to total Rab10 signal that can be diffuse (cytoplasmic) and only partially localized to vesicles (Supporting data Figure 4). The LRRK2-specific small molecule inhibitor MLi2 efficiently eliminates all pT73-Rab10 signal from these cells, suggesting the phosphorylation of Rab10 is indeed LRRK2 dependent. Several recent studies suggest chloroquine-damaged lysosomes/phagosomes may recruit LRRK2 to mediate Rab10 phosphorylation ^20^. However, our observed lack of co-localization between WT GFP(N-term)-Rab10 with the lysosomal marker LAMP1 indicates that LRRK2 might natively phosphorylate Rab10 in a different stage in phagocytic cells. To pinpoint the specific step Rab10 can be phosphorylated by LRRK2, Raw264.7 cells were co-transfected with Flag(N-term) tagged Rab10 with different phosphoinositide marker and co-labeled with pT73-Rab10 specific antibody.

**Figure 4.**
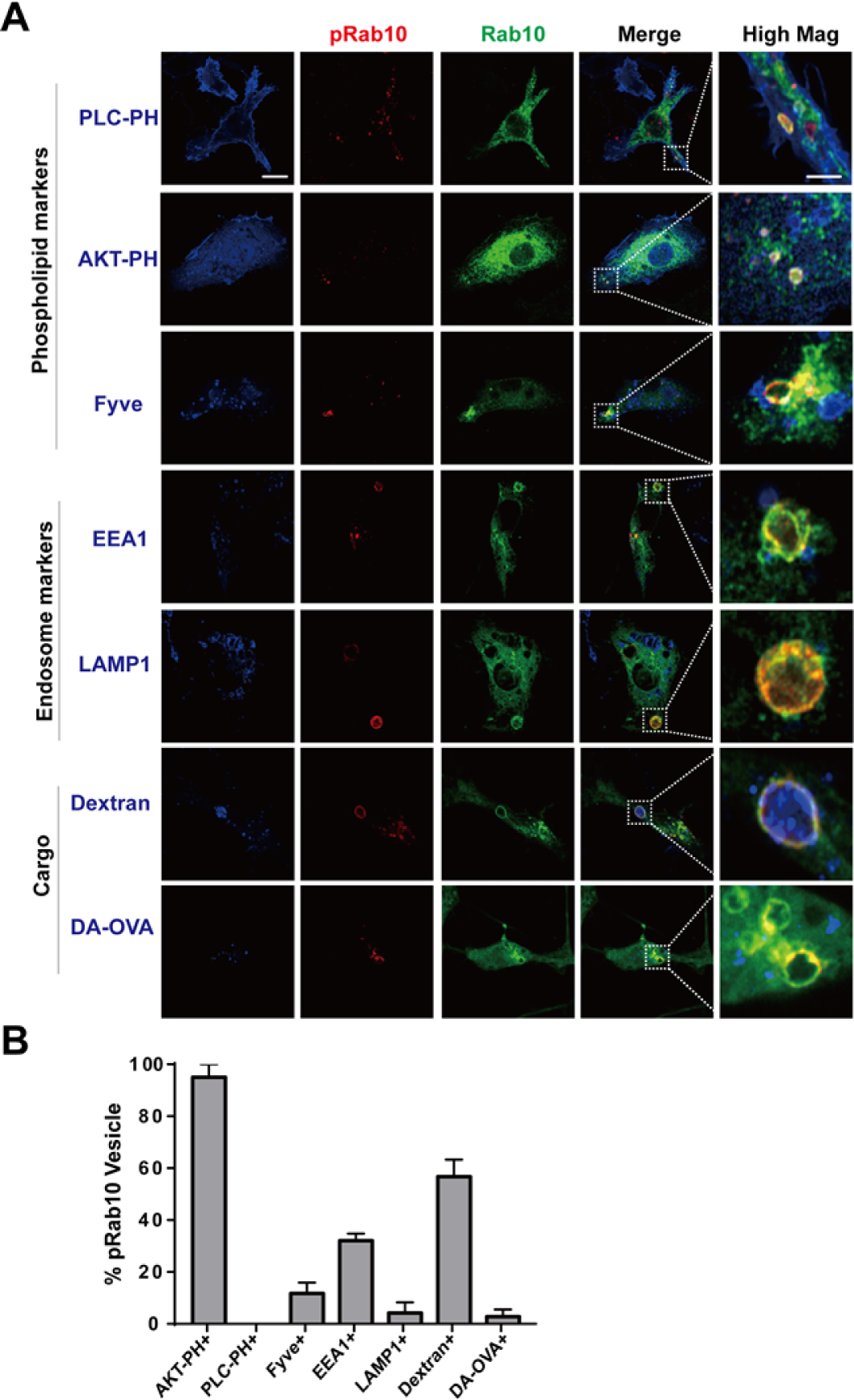
LRRK2 phosphorylates PI(3,4,5)P_3_-positive Rab10 macropinosomes in macrophage cells. **A)** To show colocalization of phosphor-Rab10 with different phosphoinositide, Raw 264.7 macrophage cells were transfected with FLAG(N-term)-Rab10 and eGFP(C-term)-AKT-PH domain(PI(3,4,5)P_3_ marker), or eGFP(C-term)-SARA-Fyve domain((PI3P marker), or eGFP(C-term)-PLC-PH domain (PI(4,5)P_2_ marker) (epifluorescence, shown in blue) followed by immune-staining with anti-Flag antibody(shown in green) and phosphor-Thr73 Rab10 specific antibody(shown in red). To show colocalization of phosphor-Rab10 with early endosome or late endosome/lysosome, Raw 264.7 macrophage cells were transfected with GFP(N-term)-Rab10 (epifluorescence, shown in green) followed by immune-staining with phosphor-Rab10 antibody (shown in red) and EEA1 or LAMP1 antibody (shown in blue). To show phosphor-Rab10 vesicles are macropinosomes, Raw 264.7 macrophage cells were transfected with GFP(N-term)-Rab10 (shown in green) and fed with 0.05mg/ml of 70kDa TRITC conjugated dextran (epifluorescence, shown in blue) for 30min followed by immune-staining with phosphor-Rab10 antibody (shown in red). To study the degradation capacity of phosphor-Rab10 vesicles, Raw 264.7 macrophage cells were transfected with FLAG(N-term)-Rab10 and fed with 0.05mg/ml of BODIPY conjugated Ovalbumin (epifluorescence, shown in blue) followed by immune-staining with anti-Flag (shown in green) and phosphor-Rab10 antibody (shown in red). Representative photomicrographs are from >30 cells analyzed for each condition from n=3 biologically independent experiments. White bounding boxes are magnified in “High Mag” panels that show individual vesicles. Scale bars represent 10 μm and 1 μm for “High Mag”. **B)** Calculated percentage of double-positive vesicles (n>30 cells counted from n=3 biologically independent experiments) are shown with error bars as ± SEM.

Similar to total Flag(N-term) tagged Rab10, >90% of pT73-Rab10 vesicles co-localize with transfected GFP(N-term) tagged AKT-PH domain PI(3,4,5)P_3_ marker, while no co-localization could be found between pT73-Rab10 and PLCδ-PH-domain (PI(4,5)P_2_ marker). Only 10% of pT73-Rab10 vesicles co-localize with SARA1-Fyve (PI3P marker, Figure 4). In contrast to previous finding showing Rab10 is predominantly phosphorylated on lysosomes ^20^, only ∼5% of pT73-Rab10 vesicles co-localize with the lysosomal marker LAMP1 (Figure 4). As expected, the majority of pT73-Rab vesicles were filled with dextran. In cells incubated with BODIPY-conjugated ovalbumin, less than 5% of pT73-Rab10 vesicles were found to be positive with BODIPY ovalbumin, suggesting these vesicles are non-degradative vesicles and consistent with the low-proportion vesicles with LAMP1 positivity (Figure 4). These data together suggest LRRK2 phosphorylates Rab10 early on PI(3,4,5)P_3_ positive macropinosomes before the vesicle can mature to and possibly merge with other EEA1/LAMP1 positive vesicles.

### LRRK2 transiently interacts with Rab10

Recent studies suggest that the phosphorylation of Rab10 is dependent on the stable membrane association of LRRK2 ^20^. Analysis of endogenous LRRK2 subcellular localization in macrophages shows a more diffusive distribution through the cytoplasm in typical non-stimulated cells in culture, with LRRK2 localization apparently unaffected by kinase inhibition in these cells (Figure 5A). In super-resolution imaging, endogenous LRRK2 protein forms transient patches of protein adjacent to pT73-Rab10-positive vesicles (Supporting data Figure 2 and Figure 5A). Biochemical fractionation experiments reveal pT73-Rab10 protein primarily localizes to triton-solubilized fractions, with LRRK2 binding to Rab10 independent of LRRK2 kinase activity (Figure 5B,C). Although pT73-Rab10 levels in these cells are exquisitely dependent on LRRK2 kinase activity, phos-tag analysis reveals pT73-Rab10 protein poorly interacts with LRRK2 compared to non-phospho Rab10 in the same triton-solubilized fraction (Figure 5,C-E). These results suggest a transient interaction supported by non-phospho LRRK2 and non-phospho Rab10 at vesicles, with complex dissociation apparently induced by LRRK2-kinase activity and phosphorylation.

**Figure 5.**
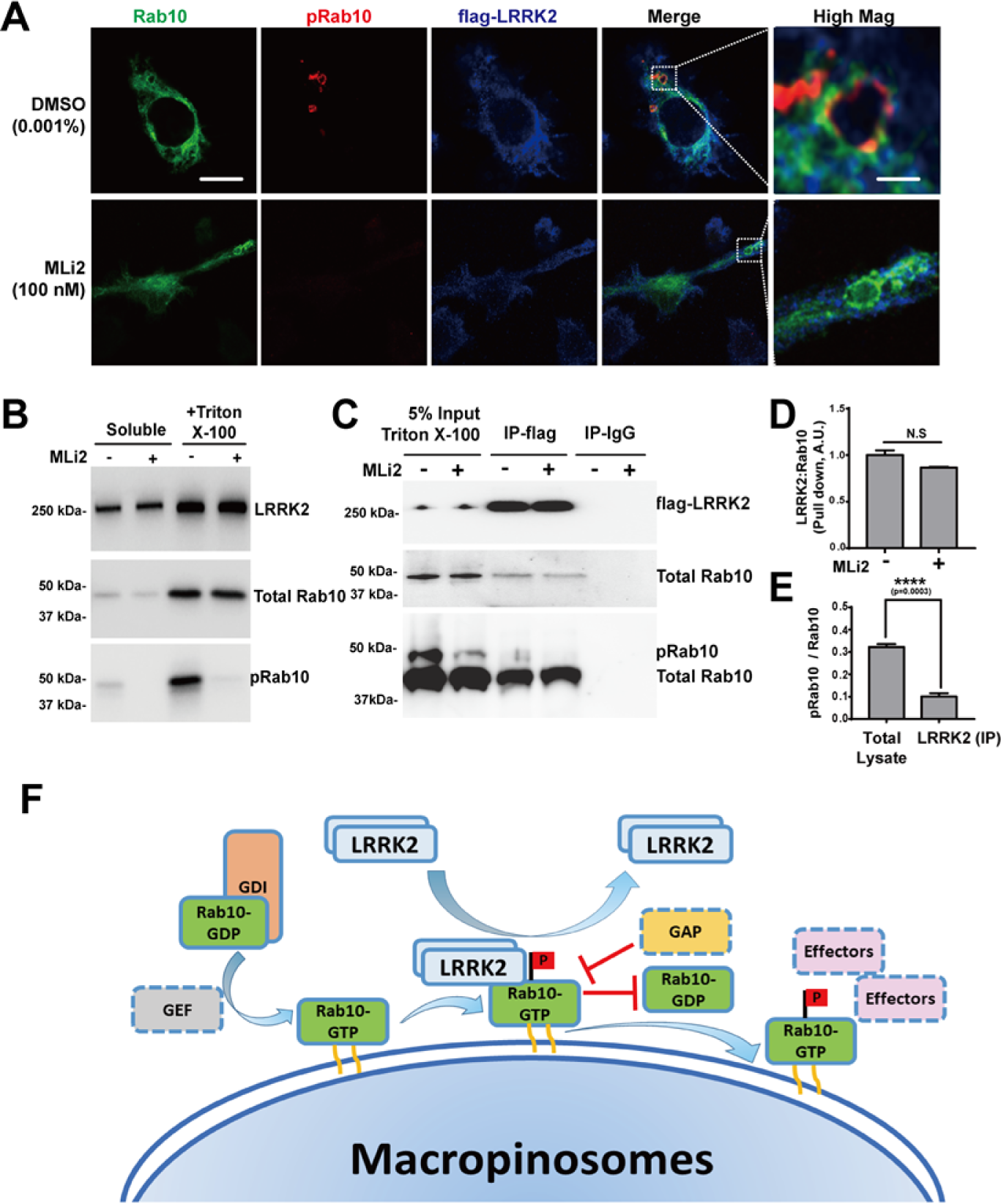
LRRK2 transiently interacts with Rab10-positive macropinosomes. **A)** Raw264.7 cells were transfected with Flag(N-term)-LRRK2 and eGFP(N-term)-Rab10 for 24 hours and treated with or without the LRRK2 kinase inhibitor MLi2 (100 nM) for 2 hours. Representative photomicrographs (from >20 images analyzed for each condition from n=3 biologically independent experiments) are shown from fixed cells with eGFP-Rab10 epifluorescence (shown as green signal), and immuno-stained for pT73-Rab10 (shown as red signal) and LRRK2 (shown as blue signal). White bounding boxes are magnified in “High Mag” panels that show individual vesicles. Scale bars represent 10 μm or 1 μm in “Hig Mag” panels. **B)** Raw 264.7 macrophage cells transfected with GFP(N-term)-Rab10 and Flag(N-term)-LRRK2 were sequentially lysed mechanically into buffer to create a ‘soluble’ protein fraction, pelleted, and then lysed into triton X-100 buffer. pT73-Rab10 was detected with a phosphor-Thr73 Rab10 specific antibody. Representative immunoblots from n=3 independent experiments are shown. **C)** The triton X-100 lysate was used as input for LRRK2 immunoprecipitation, with subsequent detection of total Rab10 and pT73-Rab10 proteins in LRRK2:Rab10 immunocomplexes. pT73-Rab10 was detected with phos-tag gel analysis. Immunoblots are representative of three independent experiments, with similar results obtained. **C)** Calculated levels of LRRK2:Rab10 with and without LRRK2 kinase inhibition, and **D)** the ratio of pT73-Rab10 to total Rab10 protein interacting with LRRK2 in input lysate and co-immunoprecipitated complex. Data are from n=3 biologically independent experiments with error bars showing ± SEM; significance was assessed by one-way ANOVA with **** representing Tukey’s post hoc test p<0.0001 or n.s. (p>0.05). **E**) Model for LRRK2 interaction with Rab protein substrates. GDP-bound Rab10 interacts with a GDI (Guanosine nucleotide dissociation inhibitor) and remain largely cytosolic. GEFs (guanine nucleotide exchange factors) facilitate the exchange of GDP for GTP and promote the macropinosome localization of Rab10. LRRK2 is transiently recruited by GTP bound active Rab10 on macropinosomes and dissociates from Rab10 once it finished phosphorylating Rab10. LRRK2-mediated phosphorylation may block the interaction of Rab10 GAPs (GTPase-activating proteins) that would otherwise facilitate the GTPase activity of Rab10 and promote Rab10 dissociation from macropinosomes ^13^. Without GAPs, Rab10 remained GTP bound and associated with macropinosomes. Therefore, LRRK2 might prolong the association of Rab10 on macropinosomes in GTP bound form. The phosphorylated Rab10 stalled on the macropinosomes could potentially affect the interaction with downstream effectors.

### LRRK2 dependent phosphorylation stalls macropinosome maturation

In native primary macrophages from mice, pT73-Rab10 vesicles represent a small proportion of Rab10 positive vesicles in the cytoplasm (Supplemental Figure 4). To further understand the consequences of LRRK2 dependent phosphorylation of Rab10, we cultured bone marrow derived macrophages from mLRRK2 WT-BAC transgenic mouse to increase the levels of LRRK2 and boost the levels of pT73-Rab (see Supporting data Figure 5). ∼80% of pT73-Rab10-macropinosomes were positive for early endosomal markers (i.e., Rab5) in contrast to total Rab10-macropinosomes (∼20%, Figure 6). Further, substantial subsets of pT73-Rab10-vesicles were positive for Rab7 or LAMP1 staining (∼30% for Rab7 and 25% Lamp1), features very rare in the total Rab10-positive macropinosome pool (∼10% for Rab7 and 5% for Lamp1 respectively, Figure 6A-C). We were unable to directly image or predict how phosphorylation affects the half-life of Rab10 macropinosomes in living cells due to the technical inability to visualize phosphorylation. However, bath-application of cells (∼20 minutes) with LRRK2 kinase inhibitor MLi2 completely ablated the pT73-Rab10-positive vesicle pool and decreased both Rab10 and Rab5-double positive vesicles to just ∼5% of total Rab10 vesicles (Figure 6 D, E). Our previous work in HEK-293 cells transfected with LRRK2 and Rab10 suggests LRRK2 phosphorylation of Rab10 may prolong Rab10 in a GTP-bound state by blocking Rab10-GAP interactions that occur near the phosphorylation site of the switch II domain ^13^. Together, these results suggest that LRRK2 mediated phosphorylation stalls Rab10 on macropinosomes to promote macropinosome maturation, mimicking GTP-locked Rab10 albeit less dramatically.

**Figure 6.**
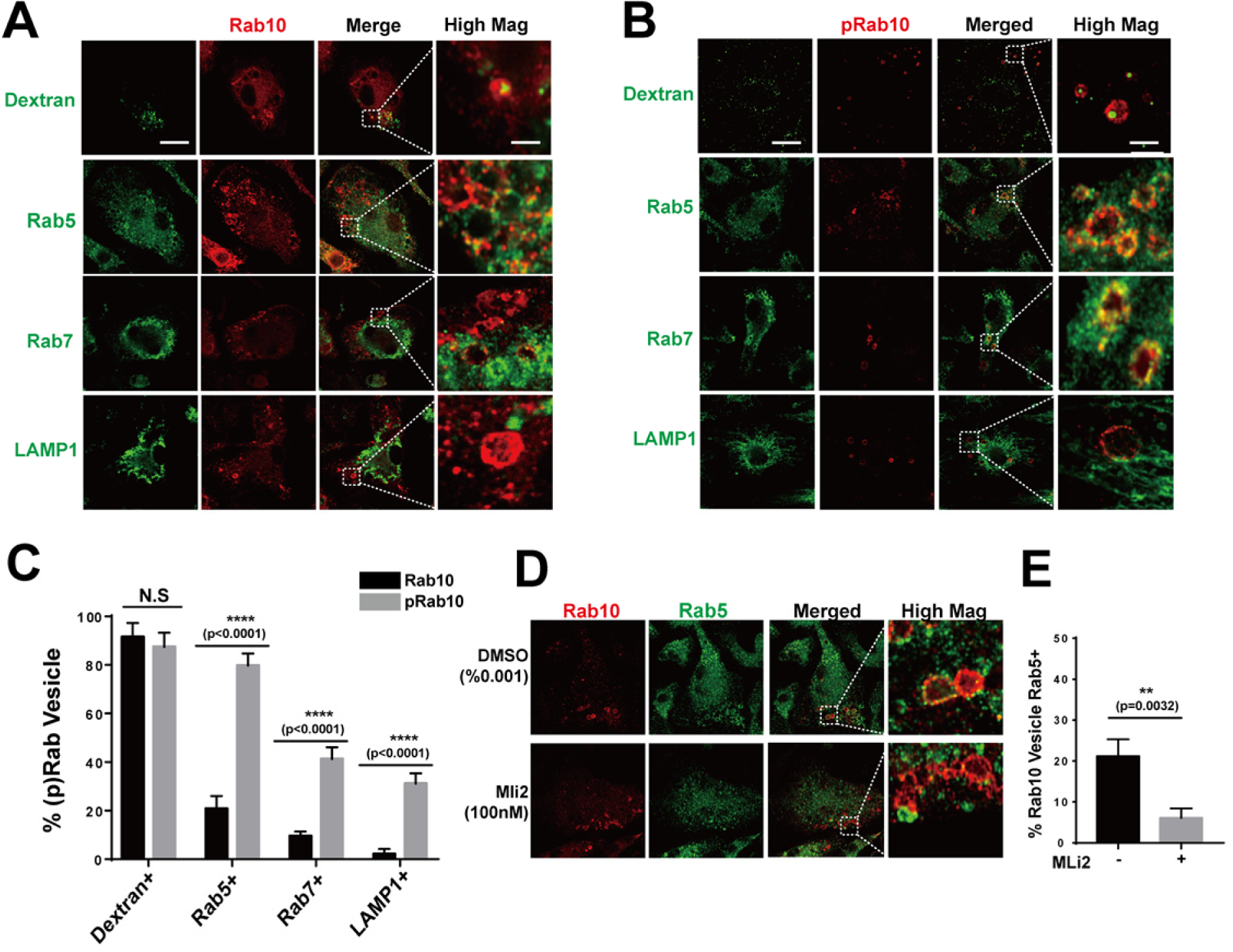
LRRK2 dependent phosphorylation stalls Rab10 on macropinosomes. **A)** Primary mouse bone marrow-derived macrophage cells (BMDM) from adult male WT-LRRK2 mBAC transgenic mice were pre-incubated with FITC-conjugated dextran (70 kDa, green signal) prior to washing, fixing, and immunostaining with total-Rab10 (shown as red signal), or **B)** pThr73-Rab10 antibody (shown as red signal). Alternatively, cells were co-stained with antibodies to Rab5, Rab7 or LAMP1 (shown as green signal in separate images). Representative photomicrographs (from >30 images analyzed for each condition from n=3 biologically independent experiments) are shown. Bounding boxes highlight vesicles in “High Mag” panels. Scale bars represent 10 μm and 1 μm for “High Mag” images. **C**) Calculated percentage of double-positive vesicles (n>30 vesicles counted from n=3 biologically independent experiments) are shown with error bars as ± SEM and significance was assessed by one-way ANOVA with Tukey’s post hoc test (** represents p< 0.01 **** represents p<0.0001 or n.s. (p>0.05) for both tests). **D)** Primary mouse bone marrow-derived macrophage cells (BMDM) from adult male WT-LRRK2 mBAC transgenic mice were treated with 0.001% DMSO or 100 nM MLi2 (i.e., >IC_90_ concentration) for 2 hours prior to fixation and immunostaining with total Rab10 or pThr73-Rab10 antibody (shown as red signal) and anti-Rab5 antibody (shown as green signal). Representative photomicrographs (from >30 images analyzed for each conditions from n=3 biologically independent experiments) are shown. Bounding boxes highlight vesicles in “High Mag” panels. Scale bars represent 10 μm and 1 μm for “High Mag” images. **E**) Calculated percentage of double-positive vesicles (n>30 vesicles counted from n=3 biologically independent experiments) are shown with error bars as ± SEM and significance was assessed by Student’s t-test with *** represents p<0.00.

Noticeably, in an attempt to further understand how LRRK2-dependent Rab10 phosphorylation alters macropinosome recycling machinery, we generated the phosphomimetic mutation T73E-Rab10 and non-phosphorylatable mutation T73A-Rab10. However, neither T73E-Rab10 or T73A-Rab10 could associate with macropinosomes (or any vesicles) in these cells (Supporting data Figure 6). Rather, mutation of the threonine 73 residue shunted Rab10 protein to aberrant perinuclear accumulations, with the T73A-Rab10 variant slightly more diffuse and noticeable mislocalization to the nucleus. As mutation of this residue is unlikely to affect GTP-binding or hydrolysis based on ours and others experiments in other cell lines, the observations here reveal the essential nature of the threonine-73 amino acid in Rab10 in mediating localization to vesicles in macrophages.

### LRRK2 mediated phosphorylation Rab10 inhibits EHBP1L1-dependent fast recycling of macropinosomes

To help identify protein co-factors that may be responsible for blocking Rab10-macropinosome vesicle recycling, we used a proteomics approach with GTP (vesicle-bound) or GDP (cytosolic)-swapped immunoprecipitated Rab10 protein combined with lysates from macrophages (Figure 7 A). This approach capitalizes on our evidence that LRRK2-mediated pT73-Rab10 partially mimics a stabilized GTP-bound Rab10. We identified only one Rab10 protein interactor exclusive to GTP-bound Rab10 that was not also bound to GDP-Rab10, the EH domain-binding protein 1-like protein 1 (EHBP1L1, Figure 7 B-D). Consistent with previous literature, pull down analysis from Raw 264.7 cells transfected with Myc-mKate2(N-term)-EHBP1L1 confirmed the specific interaction of EHBP1L1 with GTP bound but not GDP-bound Rab10 (Figure 7 C). Pull-down analysis with triton-solubilized and immobilized pT73-Rab10 and non-pT73-Rab10-beads demonstrated pT73-Rab10 weakly interacts with EHBP1L1 compared to non-phospho Rab10 (Figure 7 E, F). EHBP1L1 is well known to interact with GTP-bound Rab10 to regulate the formation of tubular recycling endosomes in Hela cells ^21, 22^. In macrophage cells transfected with eGFP(N-term)-Rab10 and Myc-mKate2(N-term)-EHBP1L1, Myc-mKate2(N-term)-EHBP1L1 prominently co-localized with Rab10 on recycling tubules. RILPL2, identified as a pT73-Rab10 specific adaptor that cannot bind non-phospho Rab10 ^14^, localized exclusively on the body of the Rab10 positive macropinosomes and never on recycling tubules (Figure 7 G,H). pT73-Rab10-vesicles, enriched with RILPL2 (Extended data figure 7), failed to form recycling tubules that are characteristic of short-lived macropinosomes, consistent with pT73-Rab10 demarcating macropinosomes that are stalled in the cytoplasm. Over-expression of EHBP1L1 blocked LRRK2-phosphorylation of Rab10 presumably through competitive access to the critical threonine-73 residue, and therefore mimicking the effects of LRRK2 kinase inhibition (Figure 7 I, J). In macrophages transfected with eGFP(N-term)-Rab10 and Myc-mKate2(N-term)-EHBP1L1, the percent of GFP(N-term)-Rab10 vesicles positive with pT73-Rab10 is significantly decreased compared to cells co-transfected with eGFP(N-term)-Rab10 and Myc-mKate2 only (Figure 7 K, L). These results favor a model whereby LRRK2 kinase activity and Rab10-phosphorylation competes with EHBP1L1-binding in mediation of macropinosome fast-recycling or maturation to EEA1-positive and finally LAMP1-positive vesicles.

**Figure 7.**
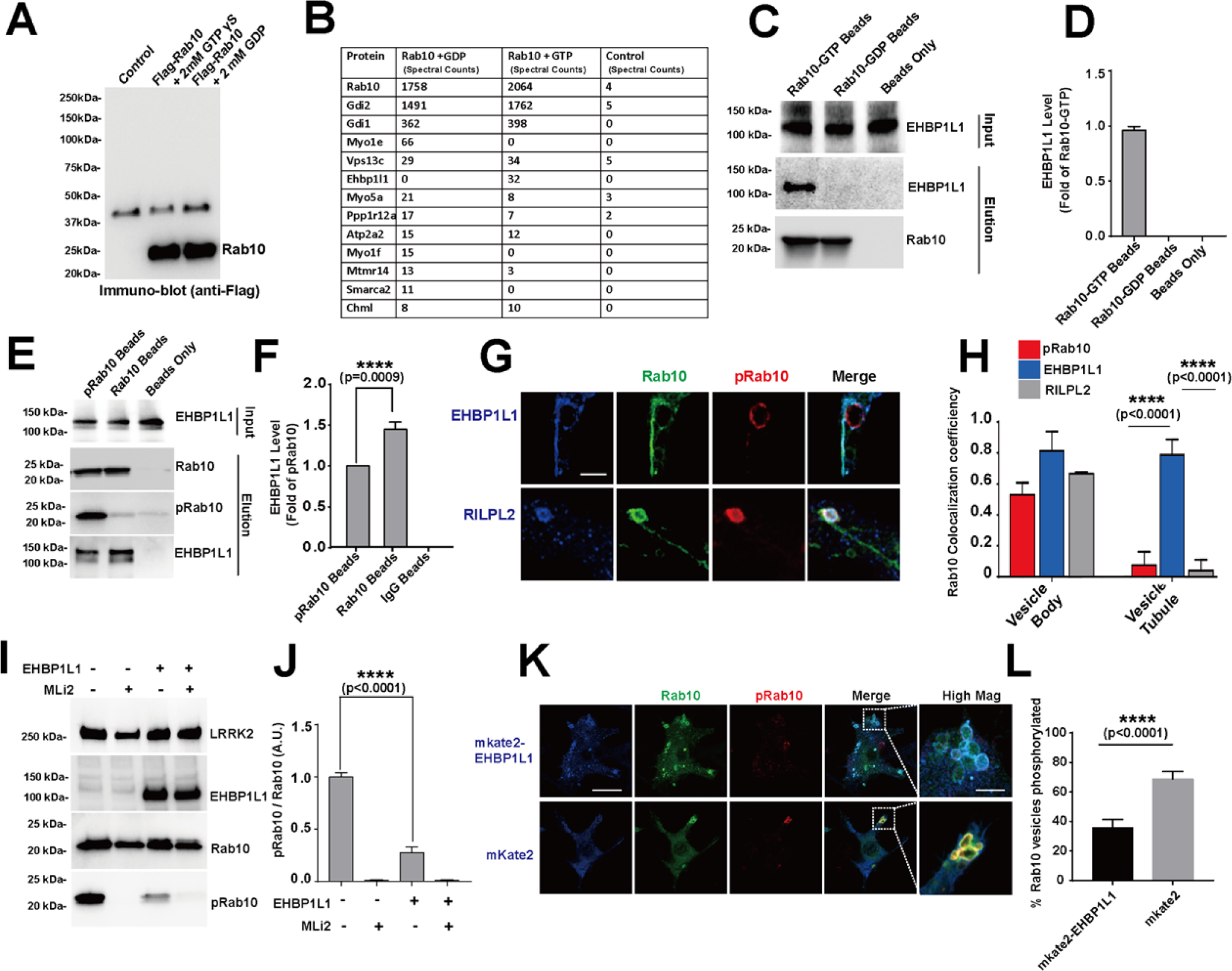
LRRK2 phosphorylation of Rab10 inhibits vesicle recycling by blocking EHBPL1 binding. **A)** Raw264.7 cells were transfected with Flag-Rab10 or control empty vector. The transfected cells were lysed in the presence of 2 mM GTPγS or 2 mM GDP. The Flag-Rab10 complex was immunoprecipitated with Flag-resin (sigma) followed by extensive washing and elution of complexes with 200 μg mL^-1^ 3xFlag-tide. (sigma). A portion of the eluted protein was analyzed by SDS-PAGE followed by immuno-blotting with anti-Flag-HRP antibody. **B)** The eluted complexes were digested with trypsin and peptides identified with Orbitrap Mass Analyzer. A representative table of peptides matching proteins with spectral counts >10 from either the GTPγS or GDP incubations, and <10 spectral counts in the control elution. **C)** Flag(N-term)-Rab10 protein is purified from HEK293 FT cells transfected with Flag(N-term)-Rab10 plasmids and immobilized on Flag-resin (sigma) in lysis buffer containing 2 mM GTPγS or 2 mM GDP. Beads are mixed with lysate of HEK293 cells transfected with Myc-mkate2(N-term)-EHBP1L1 to form Rab10:EHBP1L1 complex. The total Rab10 on the beads are detected using anti-Flag antibody. The EHBP1L1 in the lysates and immobilized on the beads are detected using anti-Myc tag antibody. Representative immunoblots from n=3 biologically independent experiments are shown. **D)** The relative EHBP1L1 pulled down level is calculated as fold of EHBP1L1 pulled down with immobilized Rab10-GTP. Data are from n=3 biologically independent experiments. **E)** HEK293 FT cells are transfected with Flag(N-term)-Rab10 and Myc(N-term)-LRRK2_R1441C_ and treated with 0.001% DMSO or 100nM MLi2. The membrane fraction of the transfected cells is enriched as described^13^ and the membrane bound Flag(N-term)-Rab10 protein is extracted with triton X-100 buffer and immobilized on Flag-tag resin followed by mixing with lysate of HEK293 cells transfected with Myc-mkate2(N-term)-EHBP1L1 to form Rab10:EHBP1L1 complex. The total Rab10 and pThr73-Rab10 on the beads are detected using anti-Flag antibody and pThr73-Rab10 antibody. The EHBP1L1 in the lysates and immobilized on the beads are detected using anti-Myc tag antibody. Representative immunoblots are shown. **F)** The relative EHBP1L1 pulled down level is calculated as fold of EHBP1L1 pulled down with immobilized Rab10 without MLi2. Data are from n=3 biologically independent experiments. **G)** Raw 264.7 cells are co-transfected with GFP(N-term)-Rab10 (epifluorescence, shown in green) and Myc-mkate2(N-term)-EHBP1L1 or Flag(C-term)-RILPL2. Cells are immune-stained with anti-Myc for EHBP1L1 or anti-Flag for RILPL2(shown in blue) and anti-pThr73 for pT73-Rab10 (shown in red). Representative photomicrographs (from >10 images analyzed for each condition from n=3 biologically independent experiments) are shown. Scale bar represents 1 μm. **H)** Pearson’s colocalization coefficient for Rab10 and pT73-Rab10, EHBP1L1 and and RILPL2 on vesicle bodies and tubular vesicles are calculated. Data are from n>10 vesicles with tubular structures from 3 independent experiments. **I)** Raw264.7 cells are co-transfected with Flag(N-term)-Rab10, Flag(N-term)-LRRK2_R1441C_ and Myc(N-term)-EHBP1L1, representative immunoblots are shown. **J)** The relative pThr73-Rab10 level is calculated as fold of cells transfected with Flag(N-term)-Rab10 and Flag(N-term)-LRRK2 only. Data are from n=3 biologically independent experiments. **K)** Raw264.7 cells are co-transfected with GFP(N-term)-Rab10 (epifluorescence, shown in green) and Myc-mkate2(N-term)-EHBP1L1 or Myc-mkate2 only (epifluorescence, shown in red). Representative photomicrographs are shown. Bounding boxes highlight vesicles in “High Mag” panels. Scale bars represent 10 μm and 1 μm for “High Mag” images. **L)** % of GFP-Rab10 vesicles positive with phosphor-Rab10 are calculated (from >20 images analyzed for each condition from n=3 biologically independent experiments). Error bars show ± SEM; significance was assessed by one-way ANOVA with * representing Tukey’s post hoc test p<0.01 or n.s. (p>0.05).

### LRRK2 dependent phosphorylation of Rab10 inhibits the fast recycling of macropinosomes

Rab10 has been previously shown to regulate TLR4 trafficking and affect LPS-stimulated pro-inflammatory cascades ^16^. However, in primary mouse bone-marrow derived macrophages, neither our Rab10 knockdown nor LRRK2 inhibitor treatment (100 nM MLi2) altered LPS-stimulated TLR4-Myd88-dependent phosphorylation of p38 or phosphorylation of AKT, proteins critical in mounting pro-inflammatory cascades in these cells (Supplemental Figure 8). Previous studies have shown that surface receptors such as G protein-coupled receptors (GPCRs), integrin and major histocompatibility complex (MHC) molecules can be internalized through macropinocytosis in phagocytes ^23–25^. Indeed, subcellular staining in mouse bone-marrow derived macrophages revealed nearly all Rab10 and pT73-Rab10 vesicles were loaded with CCR5 (GPCR), CD11b(integrin) and MHC II (Figure 8A, B). A subset of Rab10 vesicles are tabulated, with CCR5, CD11b or MHC II co-localizing with Rab10 on the tubular structures pointing toward the surface of the cells (Figure 8C). Consistent with the transfected cells, pT73-Rab10 localizes exclusively on vesicle bodies but not on the tubular structures (Figure 8 D, E) in primary macrophages. Blocking LRRK2 kinase activity in primary macrophages from mLRRK2 WT-BAC mice promotes the fast recycling of CD11b (Figure 8C-F). These data further support a model whereby LRRK2-mediated Rab10 phosphorylation inhibits the fast recycling of macropinosomes.

**Figure 8.**
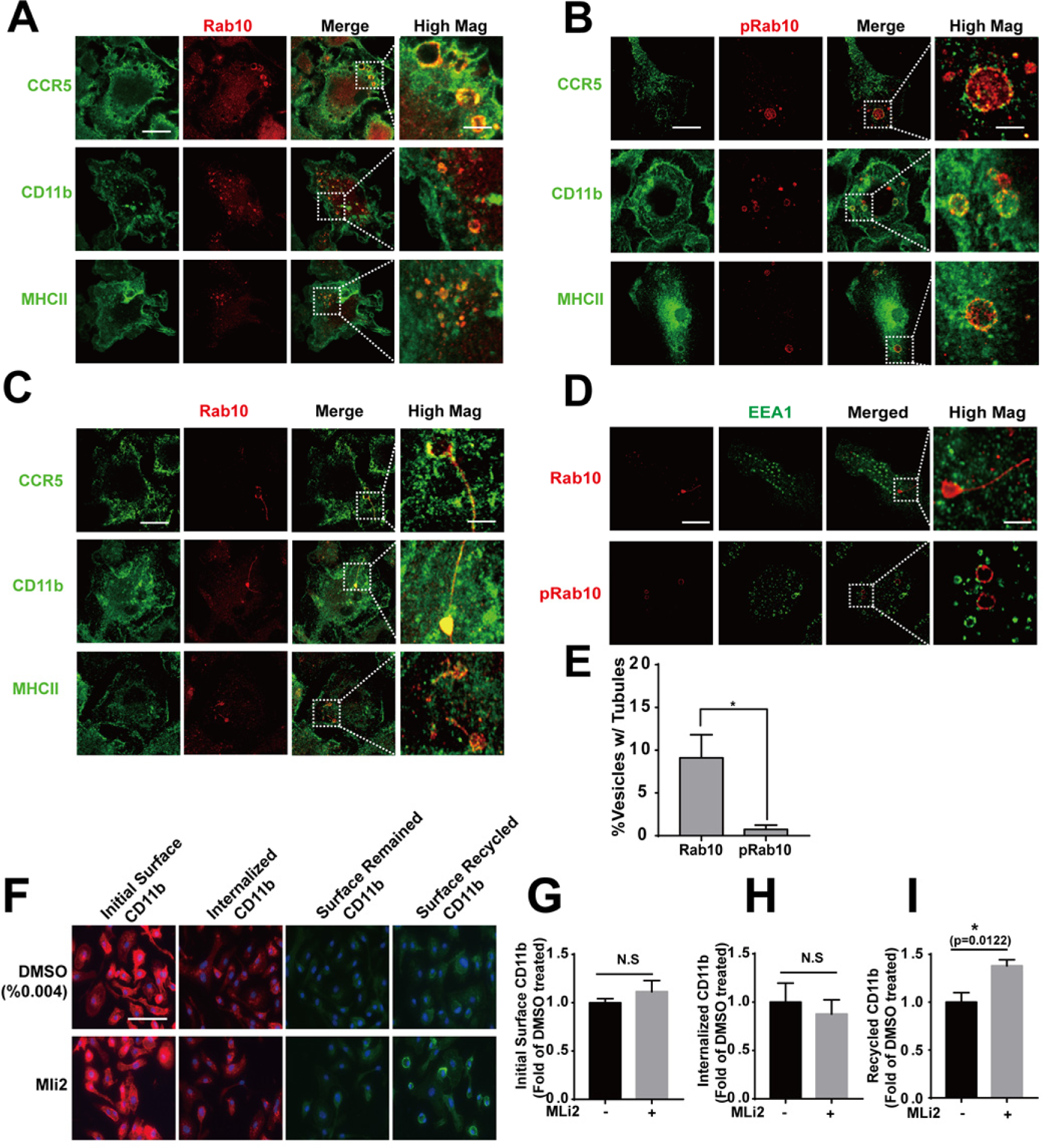
LRRK2 inhibits the fast recycling of receptors internalized through macropinocytosis. **A)** Primary mouse bone marrow-derived macrophage cells (BMDM) from adult male WT LRRK2 BAC transgenic mice are immunostained with anti-mouse CCR5, or CD11b, or MHC II antibody (green) and total Rab10 antibody or **B)** pThr73 Rab10 antibody (red) as indicated. Representative photomicrographs (from >20 images analyzed for each condition from n=3 biologically independent experiments) are shown. **C)** Representative images of CCR5, CD11b and MHC II colocalize with Rab10 on tubular structures are shown. Bounding boxes are magnified in “High Mag” panels that show individual vesicles. Scale bars represent 10 μm and 1 μm for “High Mag. **D)** Primary mouse bone marrow-derived macrophage cells (BMDM) from adult male WT LRRK2 BAC transgenic mice are immunostained with EEA1 antibody (green) and pThr73-Rab10 antibody total Rab10 antibody (red) as indicated. Representative photomicrographs (from >20 images analyzed for each condition from n=3 biologically independent experiments) are shown. Bounding boxes are magnified in “High Mag” panels that show individual vesicles. Scale bars represent 10 μm and 1 μm for “High Mag” **E)** Calculated percentage of vesicles positive with tubular structures (n>30 counted from n=3 biologically independent experiments) are shown.**F)** BMDM from adult male WT LRRK2 BAC transgenic mice are treated with %0.004 DMSO or 100nM MLi2 prior to live cell surface staining with alexa594 conjugated rat anti-mouse CD11b antibody on ice for 10 min. 3 wells of cells from each condition are fixed and imaged for initial surface CD11b (shown in red as indicated). 6 wells of cells from each condition are further incubated in 37°C to allow internalization of CD11b-antibody complex for 10 min prior to striping of surface CD11b antibody using 0.5% acetic acid on ice for 1 min. 3 of the 6 wells are fixed and surface-stained with alexa488 conjugated goat anti-rat antibody to show CD11b-antibody complex remained on the cell surface (shown in green as indicated). Internalized CD11b-antibody complex is shown in red. The other 3 wells of cells are further incubated in 37°C again for 10 min to allow recycling of internalized CD11b-antibody complex followed by surface staining with alexa488 conjugated goat anti-rat antibody to show CD11b-antibody complex recycled back to the surface. Blue represents nucleus staining. Representative images from n=3 biologically independent experiments are shown. Scale bars represent 30 μm **G)** The initial surface CD11b, **H)** internalized CD11b and **I)** recycled CD11b are calculated (from>20 images analyzed for each condition, n=3 biologically independent experiments). Error bars show ± SEM; significance was assessed by one-way ANOVA with * representing Tukey’s post hoc test p<0.05 or n.s. (p>0.05).

### Rab10 and LRRK2 potentiate CCL5-stimulated AKT-signaling

Recent studies suggest that mature macropinosomes, enriched with PI(3,4,5)P_3_, can serve as efficient signaling platforms for GPCR-induced AKT phosphorylation and subsequent signal transduction ^26, 27^. Consistent with those studies, reduction of Rab10 expression, or treatment with the LRRK2 kinase inhibitor MLi2 to block pT73-Rab10 macropinosome maturation, both potently decreased CCL5-induction of AKT phosphorylation (Figure 9 A, B). These results suggest that macropinocytosis may be a non-canonical but important pathway used by the GPCR CCR5 to mediate AKT phosphorylation in a rapid response to chemokines. LRRK2 kinase activity has been shown to augment chemotaxis both *in vivo* and *in vitro* ^28^^29^. To test whether LRRK2 dependent increase in chemotaxis response is caused by phosphorylation of Rab10 on macropinosomes, macrophages derived from mLRRK2 WT-BAC with Rab10 suppressed, or LRRK2 kinase activity blocked, failed to migrate across a Transwell based chemotaxis assay (Figure 9C,D). These data together suggest that LRRK2 mediated Rab10 phosphorylation on macropinosomes potentiates CCL5 induced chemotaxis response by promoting AKT activation.

**Figure 9.**
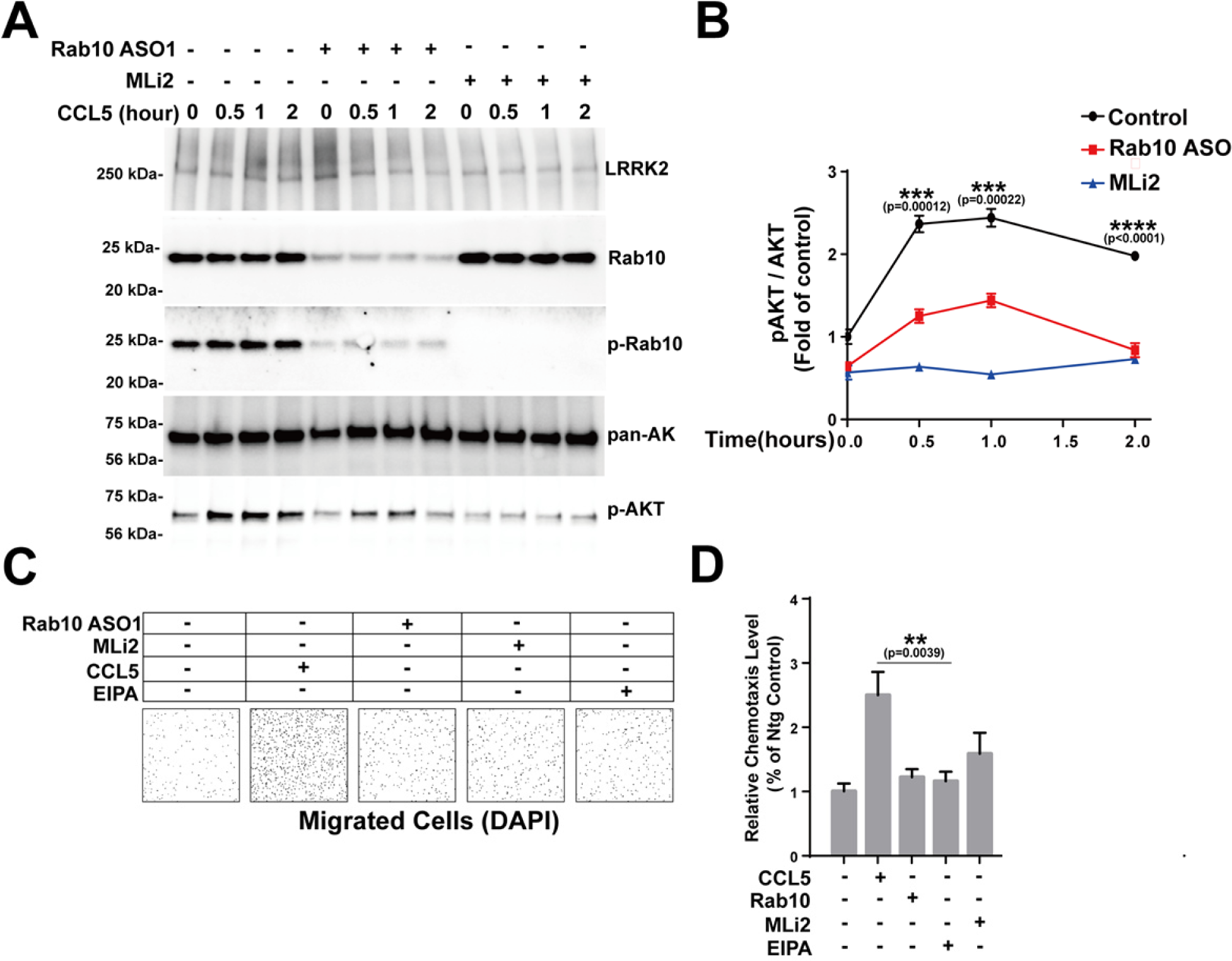
LRRK2 phosphorylation of Rab10 promotes CCL5-induced AKT activation and chemotaxis. **A)** Primary mouse bone marrow-derived macrophage cells (BMDM) from adult male WT LRRK2 BAC transgenic mice are treated with DMSO, 1 μM Rab10 ASO for 4 days or 100nM MLi2 for 12 hours before stimulated with 100nM CCL5 for indicated time points. Representative immunoblots from n=3 biologically independent experiments are shown. **B)** Relative pSer473-AKT level is normalized to total AKT and calculated as fold of cells treated without Rab10 ASO1, CCL5 or MLi2. Data are means ± SEM; significance is assessed by one-way ANOVA with Tukey’s post hoc test. ***P < 0.001, ****P < 0.0001. **C)** Primary mouse bone marrow-derived macrophage cells (BMDM) from mLRRK2 WT-BAC transgenic mice are treated with DMSO, 1 μM Rab10 ASO for 4 days, or 100nM MLi2 for 12 hours before seeded for Transwell(Sigma) chemotaxis assay using 100ng/ml CCL5 as chemoattractant. After 2 hours, the cells failing to migrate on the top layer were removed with a cotton swab. Migrated cells at the bottom of the Transwell membrane were stained with DAPI (shown as black dots) and imaged. Representative images from n=3 biologically independent experiments are shown. **D)** The relative chemotaxis level is calculated as fold of cells treated without Rab10 ASO1, CCL5, MLi2 or EIPA. Data are means ± SEM; significance is assessed by one-way ANOVA with Tukey’s post hoc test. *P < 0.05, **P < 0.01.

## DISCUSSION

Phagocytic cells (e.g., macrophages) constantly survey extracellular fluid through macropinocytosis. Early studies demonstrated ∼30% of cell volume can be internalized every hour by these macrophages, with the majority of fluid returned to the extracellular space within ten minutes ^30–32^. Mechanistic insights into macropinocytosis, a process difficult to discern from phagocytosis in practice, have been largely provided through studies of viral-entry into host cells and small molecule inhibitors ^6^. Using newly developed monoclonal antibodies to Rab10 together with live-cell imaging approaches, we initially observed Rab10 localization characteristics and dynamics that suggested possible function in macropinocytosis.

In the construction of a new model for macropinocytosis in phagocytic cells identified in this study (see Figure 10), Rab10 is recruited in formation prior to full plasma-membrane budding. Knockdown of Rab10 in a variety of phagocytic cells with constitutive macropinocytosis including human and mouse primary cells with macrophage, dendritic, or microglia-like polarizations resulted in profound impaired macropinocytosis without perturbation of phagocytosis or clathrin-dependent endocytosis of transferrin. Early macropinosomes distinct from the plasma membrane were Rab10 positive but lacked EEA1, with Rab10 interacting with EHBP1L1 in Rab10-positive tubules known to drive fast-recycling. Our data suggest that LRRK2-directed phosphorylation of the EHBP1L1 binding site (switch II effector loop) competes with EHBP1L1 binding, and consistent with recent reports, results in Rab10-binding to RILPL2 in vesicles.

**Figure 10.**
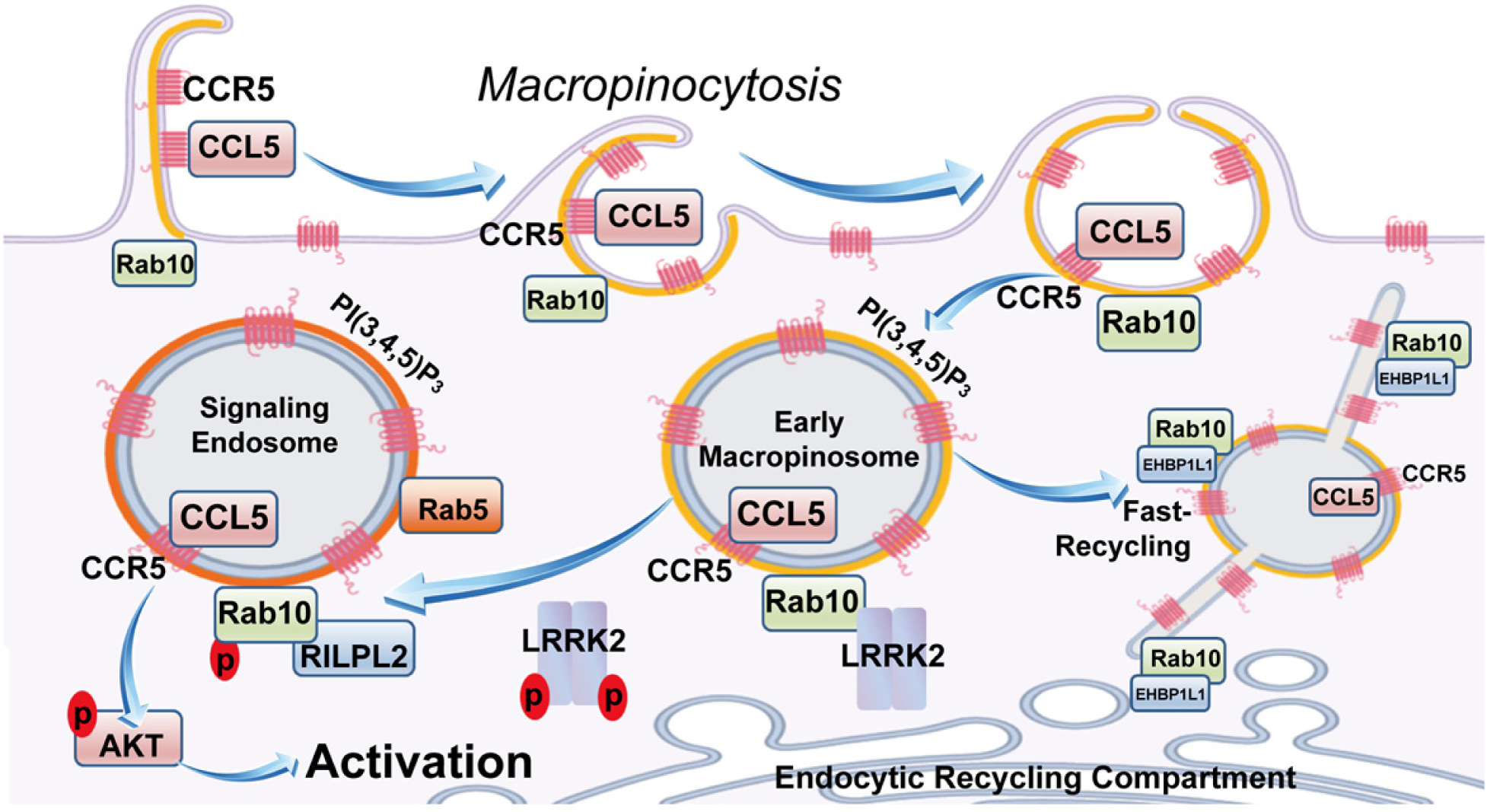
Model of the role of Rab10 and LRRK2 in regulation of macropinosome early maturation, recycling and signaling. Rab10 is initially recruited to the PI(3,4,5)P_3_ positive membrane ruffles at the early stage of macropinosome formation. During the process the ruffle closure, PI(3,4,5)P_3_ is concentrated on macropinosomes, which recruits more Rab10 to facilitate early trafficking of macropinosomes. Without LRRK2 phosphorylation, Rab10 binds EHBP1L1 and facilitate the fast recycling of surface receptors such as CCR5. LRRK2 phosphorylation of Rab10 on Thr73 inhibits the binding of Rab10 and EHBP1L1 and enhanced the binding between RILPL2 and Rab10, therefore inhibit the fast recycling of CCR5 and retained CCR5 on macropinosomes. CCR5 retained on macropinosomes can further interact with the CCL5 internalized through macropinocytosis and activates AKT signaling pathway using PI(3,4,5)P_3_ enriched macropinosomes as signaling platform.

A wealth of literature supports enrichment of CCR5 in macropinosomes that can bind HIV-1 virus for entry ^33, 34^. Consistent in our cultures, all Rab10-positive macropinosomes were also loaded with CCR5. We hypothesized that stalled macropinosomes in the cytoplasm caused by LRRK2 phosphorylation may provide an important signal transduction platform for chemokines and cytokines. CCR5 is considered a critical receptor for CCL5 (RANTES) in chemotactic responses for monocytes and T cells ^35, 36^. While T-cells lack substantial Rab10 expression, knockdown of either total Rab10 protein or pT73-Rab10 protein in macrophages blocked phosphor-AKT induction with CCL5 exposure. We propose that macropinocytosis critically complements traditional clathrin-mediated endocytosis of some GPCRs in responding to chemokines in the extracellular space through amplification of signal transduction cascades. Our initial experiments evaluating other GPCRs like CCR2 have so-far yielded ambiguous localization to macropinosomes, suggesting specificity in the pathway that requires further study. Our results may begin to clarify how macropinocytosis associated with phagocytes in the immune system use constitutive bulk-fluid uptake for surveillance and initial response to chemokines.

Both Rab10 and the protein kinase that phosphorylates Rab10, LRRK2, demonstrate selective high expression in professional phagocytic cells. In other cell types, LRRK2 phosphorylates Rab8a in mediating ciliogenesis, with Rab10 opposing this function ^37^. However, macrophages and other leukocytes are not ciliated, consistent with refined cell-type specific functions for different Rab proteins in different cells. Our previous studies in primary macrophages from mice suggest that LRRK2 kinase activity influences chemotaxis phenotypes of phagocytes in different tissues responding to a variety of immunological stimulants that include thioglycollate-induced peritonitis in the gut, over-expression of α-synuclein via rAAV2 viral transduction in the brain, or intracranial injection of LPS ^28, 38, 39^. Our results here suggest that a possible mechanism underlying these observations may involve trafficking of chemokine receptors, particularly CCR5 known to be important in some of these responses. Our results suggest LRRK2 kinase activity may serve to stabilize a fraction of macropinosome-originating signaling endosomes to bolster chemokine-receptor signaling and thus amplify the effects of chemokines like CCL5 in activating and mobilizing immune cells responding to stimuli. With both Rab10 and LRRK2 genetically linked to proteinopathies that include AD and PD, respectively, and clear roles for phagocytic immune cells in the uptake and clearance of misfolded proteins, we further speculate that the pathway discovered here may explain in part how LRRK2 and Rab10 contribute to neurodegeneration in immune cells.

## METHODS

### Plasmids and constructs

pCDNA3.1-GFP-Rab10 and pCDNA3.1-Flag-Rab10 were generated by insertion of synthesized Rab10 cDNA into pCDNA3.1+N-EGFP plasmid or pCDNA3.1+/N-DYK plasmid from Genscript as described ^13^. mRuby-Rab5 is kindly provided by Dr. Laura Volpicelli-Daley from University of Alabama at Birmingham. GFP-Fyve is generated by insertion of synthesized SARA1 Fyve domain into pCDNA3.1+N-EGFP plasmid from Genscript as described ^40^. GFP-AKT-PH (plasmid# 21218) and GFP-PLC δ –PH (plasmid # 51407) plasmids are acquired from Addgene. pCDNA3.1-Flag-LRRK2_R1441C_ is generated as previously described^41^. pCDNA3.1-RILPL2-Flag is purchase from Genscript(OHu17095). pCDNA3.1-Myc-mKate2-EHBP1L1 is generated by insertion of synthesized mKate2 and EHBP1L1(OMu05300C, Genscript) into pcDNA3.1+N-MYC plasmid from Genscript.

### Cell Culture

Mouse bone marrow derived macrophages are generated by culturing the mouse bone marrow cells collected from 3-5 months old mouse in 10% Fetal bovine serum and DMEM supplemented with 20 ng/ml M-CSF. Mouse bone marrow derived dendritic cells are generated by culturing the mouse bone marrow cells in 10% Fetal bovine serum and DMEM supplemented with 20 ng/ml GM-CSF. Mouse primary microglia is collected from new born mouse and cultured as previously described ^42^.

Human Monocyte were acquired from frozen PBMCs with STEMCELL EasySep Human Monocyte Enrichment kit (STEMCELL TECHNOLOGIES). Purified monocytes were cultured in RPMI-1640 Glutamax medium (Invitrogen) supplemented with 10% fetal bovine serum (FBS; Atlanta Biological), penicillin/streptomycin (Lonza) and Fungizone (2.5 ug/ml; Life Technologies). GM-CSF (all cytokines come from Peprotech, 10 ng/ml) or M-CSF (20 ng/ml) were supplemented into the monocyte culture at day 5 to induce dendritic-like cells and macrophage-like cells respectively. MDMi cells were induced as previous protocol described ^43^, briefly, monocytes were cultured in serum free RPMI-1640 Glutamax medium (Invitrogen) for 10 days before supplemented with GM-CSF 10 ng/ml, M-CSF 10 ng/ml, NFG-β 10 ng/ml, CCL2 100ng/ml, and IL-34 100 ng/ml.

Raw 264.7 cells (TIB-71, ATCC) were cultured in 10% Fetal bovine serum and DMEM. Raw 264.7 cells are collected at 60% confluency for transfection. Transfection was performed using Neon® Transfection System (Thermo Fisher Scientific) following the manufactory’s protocol. The cells are either collected for immunoblots or fixed for microscopy 12 hours after transfection.

### ASO synthesis, screening and lead identification

Synthesis and purification of all chemically purified ASOs were performed as previously described (Seth et al., J Org. Chem 2010). Approximately 500 ASOs were designed against the full mouse *Rab10* gene. ASOs were screened in days in vitro 7 (DIV7) primary mixed cortical neurons derived from embryonic day 16 C67BL/6N mice at 7 µM. ASOs were applied to culture medium for 24h, after which cells were harvested for RNA extraction and mouse *Rab10* mRNA was quantified by RT-PCR using the forward primer, reverse primer, and probe. *Rab10* ASO 1 and 2 (Table 1) were identified among the dose-responsive ASOs in the follow-up dose-response confirmation treatment.

### Immunoblotting

Protein lysates were analyzed using SDS-PAGE followed by transfer to PVDF membranes for immunoblotting with indicated primary antibodies and HRP-conjugated secondary antibodies. Signals developed with Classico ECL reagent (Millipore) was digitally detected using Chemidoc Touch (BioRad). Intensities of the indicated bands are calculated with ImageLab software. The following antibodies were used: N241A/34 anti-LRRK2 (Antibodies Inc), phospho-T73-Rab10(MJF-R21), anti-eGFP antibody (ab6673, Abcam), total Rab10 antibody (Cell signaling), anti-FLAG M2 (Sigma), anti-Myc antibody (ab32 Abcam), p38 antibody (8690, cell signaling), phospho-p38 antibody (4511, cell signaling), IRF3 antibody(4302, cell signaling), phospho-IRF3 antibody(4947 cell signaling), total AKT antibody(4691), phospho-AKT antibody(4060), β-actin (sc-47778 HRP, Santa Cruz).

### Immunoprecipitation

Raw264.7 cells transfected with pCDNA3.1-Flag-Rab10 are lysed with lysis buffer containing 150 mM NaCl, 50 mM Tris-HCl, pH 7.4, 10mM MgCl_2_, 0.05% Triton-100, 1x PhosSTOP and Protease inhibitor cocktails (Roche) followed by ultracentrifugation at 150, 000 x g for 20 min. The supernatant was collected and mixed with anti-Flag resin (M8823, sigma) for 12 hours in 4C. The beads are washed 5 times with lysis buffer and eluted with lysis buffer supplemented with 100µg/ml 3X FLAG® Peptide (F4799, sigma). The eluted lysates are analyzed using SDS-PAGE or mass spectrometer.

### Mass spectrometry

Samples were reduced with 10 mM dithiolthreitol for 30 min at 80C and alkylated with 20 mM iodoacetamide for 30 min at room temperature. Next, they were supplemented with a final concentration of 1.2% phosphoric acid and 1084 µL of S-Trap (Protifi) binding buffer (90% MeOH/100mM TEAB). Proteins were trapped on the S-Trap, digested using 20 ng/µl sequencing grade trypsin (Promega) for 1 hr at 47C, and eluted using 50 mM TEAB, followed by 0.2% FA, and lastly using 50% ACN/0.2% FA. All samples were then lyophilized to dryness and resuspended in 12 µL 1%TFA/2% acetonitrile containing 12.5 fmol/µL yeast alcohol dehydrogenase (ADH_YEAST).

Quantitative LC/MS/MS was performed on 4 µL of each sample, using a nanoAcquity UPLC system (Waters Corp) coupled to a Thermo Orbitrap Fusion Lumos high resolution accurate mass tandem mass spectrometer (Thermo) via a nanoelectrospray ionization source. Briefly, the sample was first trapped on a Symmetry C18 20 mm × 180 µm trapping column (5 μl/min at 99.9/0.1 v/v water/acetonitrile), after which the analytical separation was performed using a 1.8 µm Acquity HSS T3 C18 75 µm × 250 mm column (Waters Corp.) with a 90-min linear gradient of 5 to 30% acetonitrile with 0.1% formic acid at a flow rate of 400 nanoliters/minute (nL/min) with a column temperature of 55C. Data collection on the Fusion Lumos mass spectrometer was performed in a data-dependent acquisition (DDA) mode of acquisition with a r=120,000 (@ m/z 200) full MS scan from m/z 375 – 1500 with a target AGC value of 2e5 ions. MS/MS scans were acquired at Rapid scan rate (Ion Trap) with an AGC target of 5e3 ions and a max injection time of 100 ms. The total cycle time for MS and MS/MS scans was 2 sec. A 20s dynamic exclusion was employed to increase depth of coverage. The total analysis cycle time for each sample injection was approximately 2 hours.

The MS/MS data was searched against the SwissProt M. musculus database (downloaded in Apr 2017) and an equal number of reversed-sequence “decoys” for false discovery rate determination. Mascot Distiller and Mascot Server (v 2.5, Matrix Sciences) were utilized to produce fragment ion spectra and to perform the database searches. Database search parameters included fixed modification on Cys (carbamidomethyl) and variable modifications on Meth (oxidation) and Asn and Gln (deamidation).

### Phos-tag gel analysis

PT73-Rab10 co-immunoprecipitated with LRRK2 is analyzed using SDS-PAGE gel supplemented with 100 µM MnCl2 and 50 µM of Phos-Tag reagent (Wako Chemicals) as previously described ^13 44^. Gels are transferred to PVDF membrane followed by immunostaining using anti-Flag antibody.

### Endocytosis assays

Dextran uptake assay is done by mixing of 0.1mg/ml of fluorescein labeled 70kDa Dextran (D1822, invitrogen) or tetramethylrhodamine labeled 70kDa dextran (D1818, Invitrogen) with indicated cells in 48 wells for 30mins. Phagocytosis assay is done by mixing of 0.1mg/ml of mouse IgG labeled Fluoresbrite® 641 Carboxylate Microspheres (polysciences, 17797-1) with indicated cells for 30mins. Transferrin uptake assay is done by mixing of 0.1mg/ml of alexa-488 labeled transferrin (T13342, Invitrogen) with indicated cells for 30min. Macropinosome maturation assay is done by mixing of 0.1mg/ml of DQ-conjugated ovalbumin (D12053, Invitrogen) with indicated cells for indicated time points. Phagosome maturation assay is done by mixing DQ-ovalbumin conjugated Fluoresbrite® 641 Carboxylate Microspheres with indicated cells for 30mins. Cells are washed 3 times with phosphate-buffered saline fixed with 4% paraformaldehyde for 20mins before imaging.

### CD11b recycling assays

BMDM from adult male WT LRRK2 BAC transgenic mice are treated with %0.004 DMSO or 100nM MLi2 prior to live cell surface staining with alexa594 conjugated rat anti-mouse CD11b antibody on ice for 10 min. A total of 9 wells of cells for each condition are treated as described below-3 wells of cells from each condition are fixed with 4% paraformaldehyde and imaged for alexa594 fluorescence to indicate initial surface CD11b. 6 wells of cells from each condition are further incubated in 37°C to allow internalization of CD11b-antibody complex for 10 min prior to striping of surface CD11b antibody using 0.5% acetic acid, 0.5M NaCL pH 3.0 on ice for 1 min. Cells are washed 5 times with DMEM media to equilibrate the pH back to 7.4. 3 of the 6 wells of cells are fixed and surface-stained with alexa488 conjugated goat anti-rat secondary antibody. The alexa594 fluorescence images are recorded to indicate the amount of CD11b-antibody complex internalized. The alexa488 fluorescence images are recorded to indicate the amount of CD11b-antibody remained on the surface after acetic acid stripping. The last 3 wells of cells are incubated in 37°C again for 10 min to allow recycling of internalized CD11b-antibody complex followed by surface staining with alexa488 conjugated goat anti-rat antibody to show CD11b-antibody complex recycled back to the surface.

### Chemotaxis assays

Primary mouse bone marrow-derived macrophage cells (BMDM) from adult male Ntg or WT LRRK2 BAC transgenic mice are treated with DMSO, 1 μM Rab10 ASO for 4 days or 100nM MLi2 for 12 hours before lifted with Accutase (Sigma, A6964-100ML) and seeded on the upper chamber of Transwell(Corning Costar, CLS3464-12EA). Cells are incubated for 2 hour in 37°C before CCL5 (eBioscience, 34-8385-64) was added in the bottom chamber. After 2 hours, the unmigrated cells on the upper surface of the Transwell membrane was removed using cotton swab. The migrated cells at the bottom surface of the Transwell membrane were fixed with 4% paraformaldehyde and stained with DAPI (shown as black dots) and imaged.

### Immuno-fluorescence staining

Cells are fixed with 4% paraformaldehyde, washed 3 time with phosphate-buffered saline(PBS), permeabilized with PBS supplemented with 0.1% saponin, blocked with 5% goat serum and immune-stained for 12 hours with primary antibodies followed by 12hours staining with secondary antibody. Horchest is used for nucleus staining. For confocal microscopy or airyscan microscopy, cell culture coverslips are mounted on glass slides with ProLong Gold antifade mountant (Thermo Fisher Scientific). The following antibodies were used: N241A/34 anti-LRRK2 (Antibodies Inc), phospho-T73-Rab10(MJF-R21-22-5), total RAB10 antibody (MJF-R23), anti-eGFP antibody (ab6673, Abcam), anti-FLAG M2 (Sigma), anti-Myc antibody (ab32 Abcam), EEA1(sc-365652, santa cruz), LAMP1(sc-19992), CD11b(14-0112-82, Thermo Fisher Scientific), TLR4 (ab22048, Abcam), CCR5(17-1951-80 Thermo Fisher Scientific).

### Widefield and super-resolution microscopy

For endocytosis assays, cells in 48 wells are imaged using Keyence bz-x700 microscope with 20x objective lens. For live cell imaging, cells cultured in 35mm glass bottom plates are imaged using Keyence microscope with 63x Oil-Immersion Objective lens in live cell imaging chamber at 37C supplied with 5% CO_2_. For high-resolution imaging, cells mounted on glass slides are imaged using Zeiss 880 Airyscan Inverted Confocal microscope with 63x Oil-Immersion Objective lens. The Airyscan images are processed using Zen 3.0 software from Zeiss.

### Mice

All the mice (3-5 months of age, males and females) used in this study were bred at the University of Alabama at Birmingham (UAB). All protocols used in this study were approved by UAB Animal Care and Use Committee that is fully accredited by the AAALAC. Mouse FLAG-WT LRRK2 BAC (B6.Cg-Tg(Lrrk2)6Yue/J), LRRK2 knock out (KO) mice and non-transgenic (nTg) littermate controls (C57bL/6J) including their genotyping strategies have been described elsewhere (53-55). Quantitative PCR was used to determine the copy number of the BAC transgene from these transgenic mice and only the mice with 20-30 copies of the BAC transgene were used in this study.

### Statistical analyses

GraphPad Prism 6 (La Jolla, CA 92037) was used to perform all the statistical analyses: 1 way-ANOVA (with Tukey’s multiple comparison test), 2 way-ANOVA (with Tukey’s multiple comparison test) and student’s t-test). A p value of <0.05 was considered significant for comparing groups.

## SUPPLEMENTAL FIGURES

**Supplemental Figure 1.**
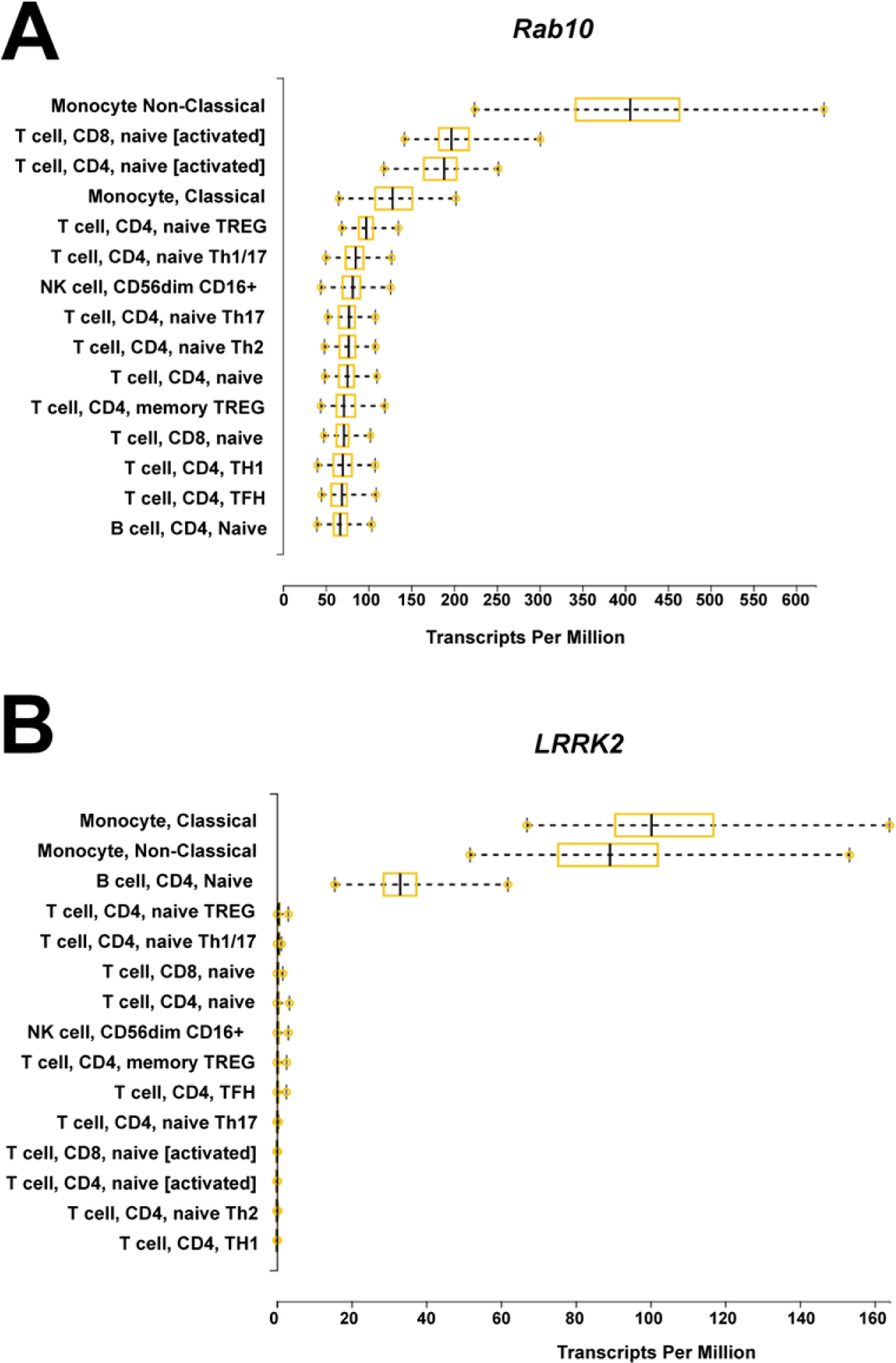
Rab10 and LRRK2 selective co-expression in monocytes. **A)** Expression profiles of Rab10 and **B)** LRRK2 in different immune cells. Results are adapted from DICE (Database of Immune Cell Expression, Expression quantitative trait loci eQTLs, and Epigenomics, https://dice-database.org/)

**Supplemental Figure 2.**
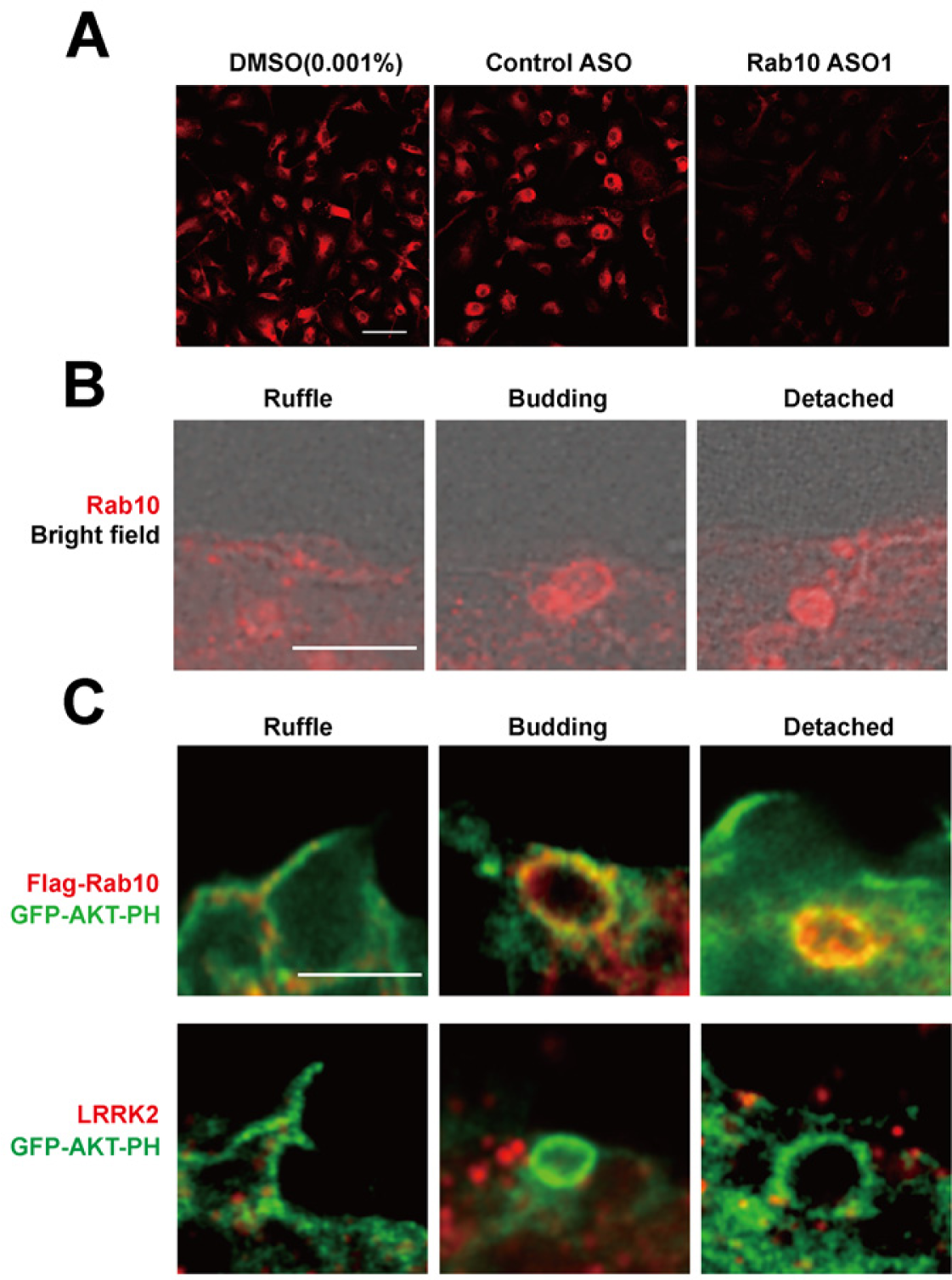
Rab10-positive vesicles emergent from ruffled plasma membrane in macrophages. **A)** Mouse bone marrow derived macrophages (BMDMs) treated with DMSO, control ASO or ASO1 were fixed and immune-stained with anti-total-Rab10 antibody (shown in red). Representative confocal images are shown. Scale bar represents 50 μm. **B)** Mouse bone marrow derived macrophages (BMDMs) were fixed and immuno-stained with anti-total-Rab10 antibody (shown in red). Plasma membrane was imaged with wide-field fluorescence microscopy overlaid with phase contrast (shown in grey). Representative images are shown. Scale bar represents 2 μm **C)** Raw264.7 cells were co-transfected with Flag(N-term)-Rab10 or LRRK2, with eGFP-tagged AKT-PH domain. Cells were fixed and immuno-stained with Flag-tag antibody, LRRK2 (shown in red), with epifluorescence (shown in green). Representative photomicrographs shown. Scale bar represents 1 μm.

**Supplemental Figure 3.**
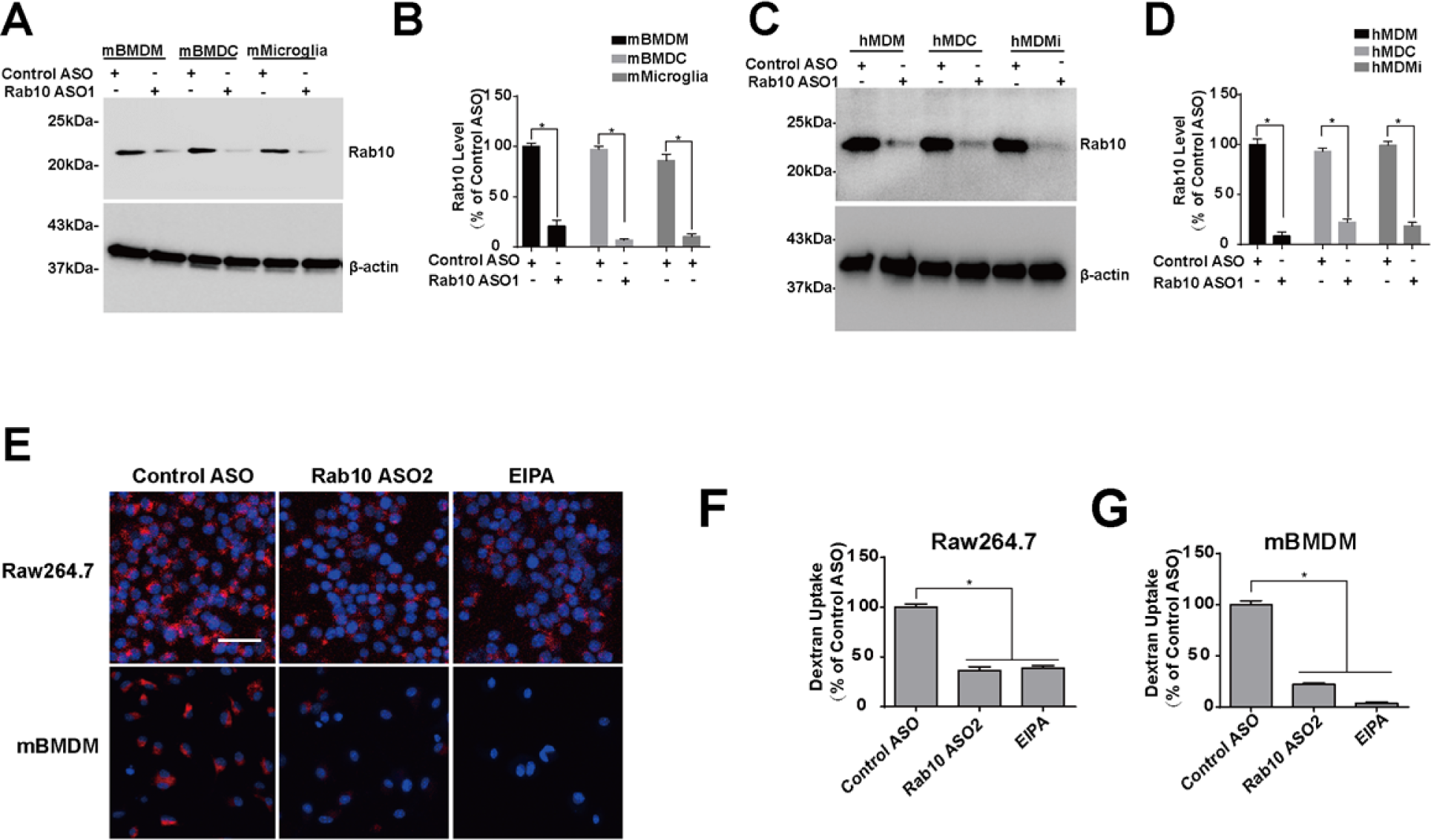
Rab10 expression is required for macropinocytosis in human and mouse phagocytic cells. **A)** Primary mouse bone marrow-derived macrophage cells (BMDMs) from adult male C57BL/6J mice were differentiated to a monocytic lineage (mBMDM), dendritic lineage (mBMDC), or microglia-like lineage (mMicroglia). Cells were treated with 1 μM control ASO or Rab10 ASO1 for four days before harvesting protein lysate and immunoblotting. Representative immunoblots from three biologically independent experiments are shown. **B)** Relative Rab10 level is calculated as % of mBMDM cells treated with control ASO. **C)** Human monocytes purified from human venous blood from healthy adult males were were differentiated to a monocytic lineage (hMDM), dendritic lineage (hMDC), or microglia-like lineage (hMDMi). Cells were treated with 1 μM control ASO or Rab10 ASO1 for four days before harvesting protein lysate and immunoblotting. Representative immunoblots from three biologically independent experiments are shown. **D)** Relative Rab10 level is calculated as % of hMDM cells treated with control ASO. **E)** Raw264.7 cells and mBMDM cells derived from C57BL/6J mice were treated with control ASO or Rab10 ASO2 for 4 days, or 50 μM EIPA (5-[N-ethyl-N-isopropyl]amiloride) for 20 min, prior to incubation for 60 min with fluorescent (tetramethylrhodamine, TRITC)-labeled dextran (70 kDa, shown as red signal). Representative photomicrographs (from >20 images analyzed for each conditions from n=3 biologically independent experiments) are shown from cells extensively washed, fixed, and stained with DAPI (shown as blue signal). **F, G)** Relative fluorescent signals were calculated as a percent of control-ASO treated cells. Scale bars show 50 μm. Column graphs show group means with error bars as ± SEM; significance was assessed by one-way ANOVA with * representing Tukey’s post hoc test p<0.01 or n.s. (p>0.05).

**Supplemental Figure 4.**
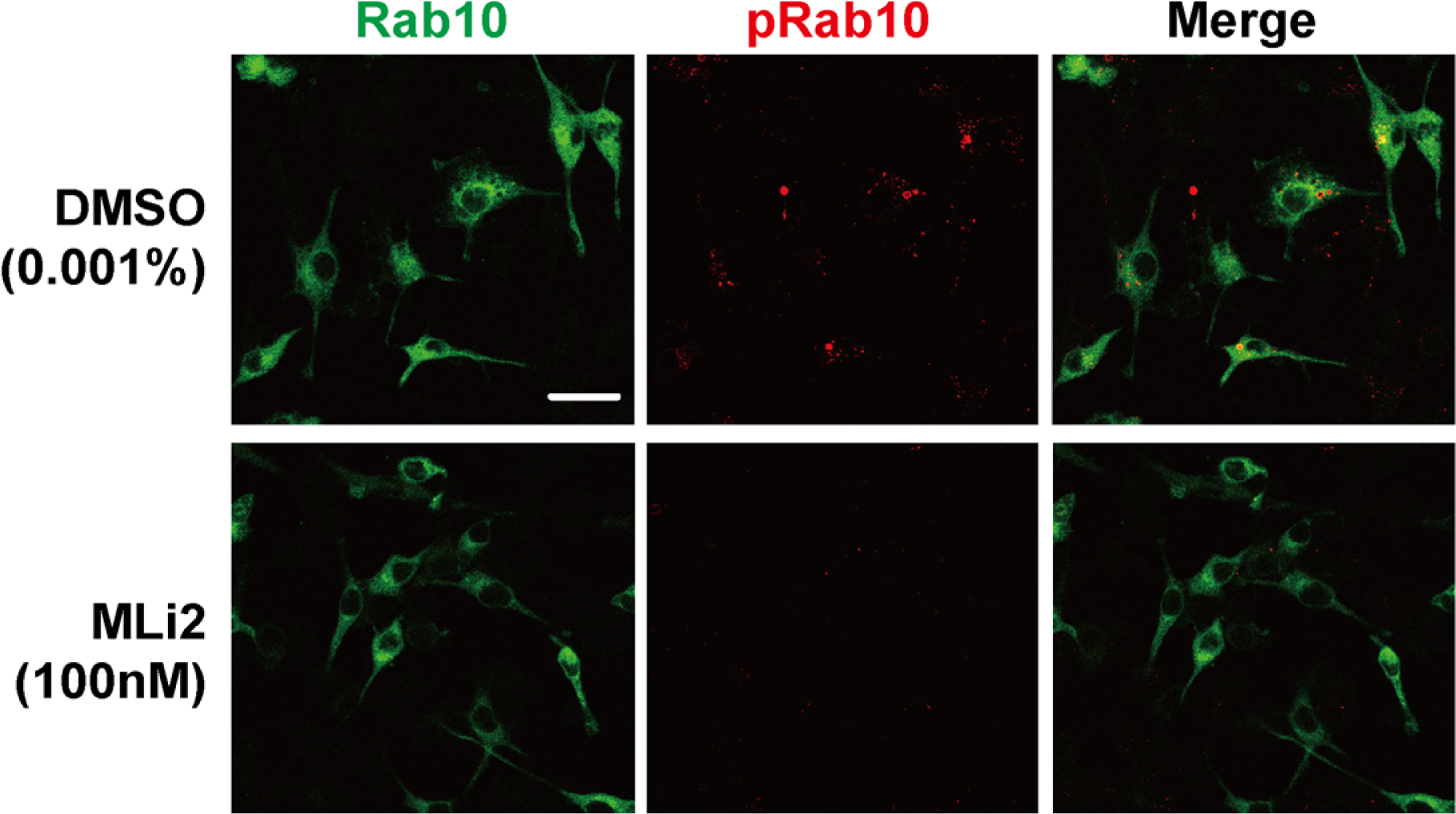
Rab10 expression is required for macropinocytosis in human and mouse phagocytic cells. D) Raw264.7 cells were transfected with eGFP(N-term)-WT Rab10 (show as green signal) and treated with 0.001% DMSO or 100 nM MLi2 (i.e., >IC_90_ concentration) for 2 hours prior to fixation and immunostaining with pThr73-Rab10 antibody (shown as red signal). Representative photomicrographs (from >20 images analyzed for each conditions from n=3 biologically independent experiments) are shown. Scale bars indicate 20 μm.

**Supplemental Figure 5.**
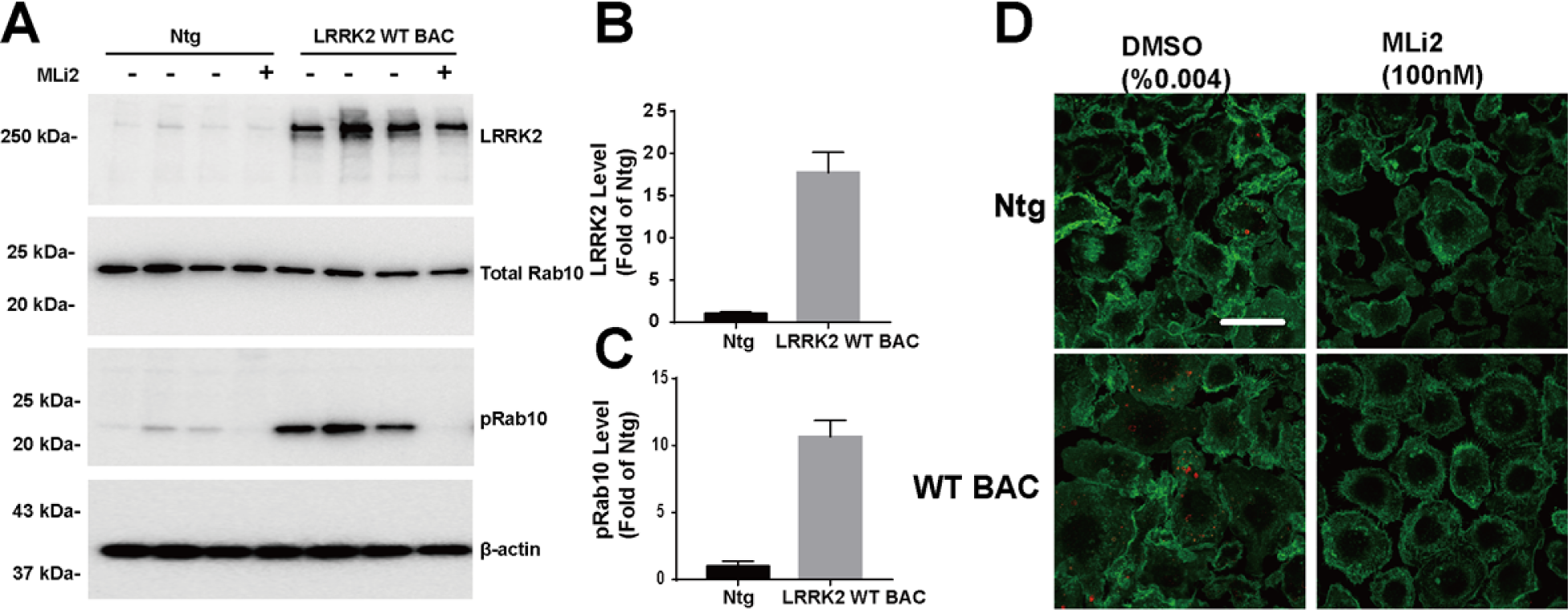
LRRK2 WT BAC express high level of LRRK2 and phosphor-Rab10. **A)** Primary mouse bone marrow-derived macrophage cells (BMDM) from adult male Ntg or LRRK2 WT BAC transgenic mice are treated with DMSO or 100nM Mli2 for 2 hours. Representative immunoblots of cells derived from 3 different animals for each genotype are shown. **B)** Relative LRRK2 or **C)** pT73-Rab10 level is calculated as fold of Ntg cells treated with DMSO. **D)** Primary mouse bone marrow-derived macrophage cells (BMDM) from adult male Ntg or LRRK2 WT BAC transgenic mice are treated with DMSO or 100nM Mli2 for 2 hours and immune-stanined with CD11b (shown in green) and pThr73 Rab10(shown in red). Representative images from N>10 images are shown. Scale bars indicate 50 μm.

**Supplemental Figure 6.**
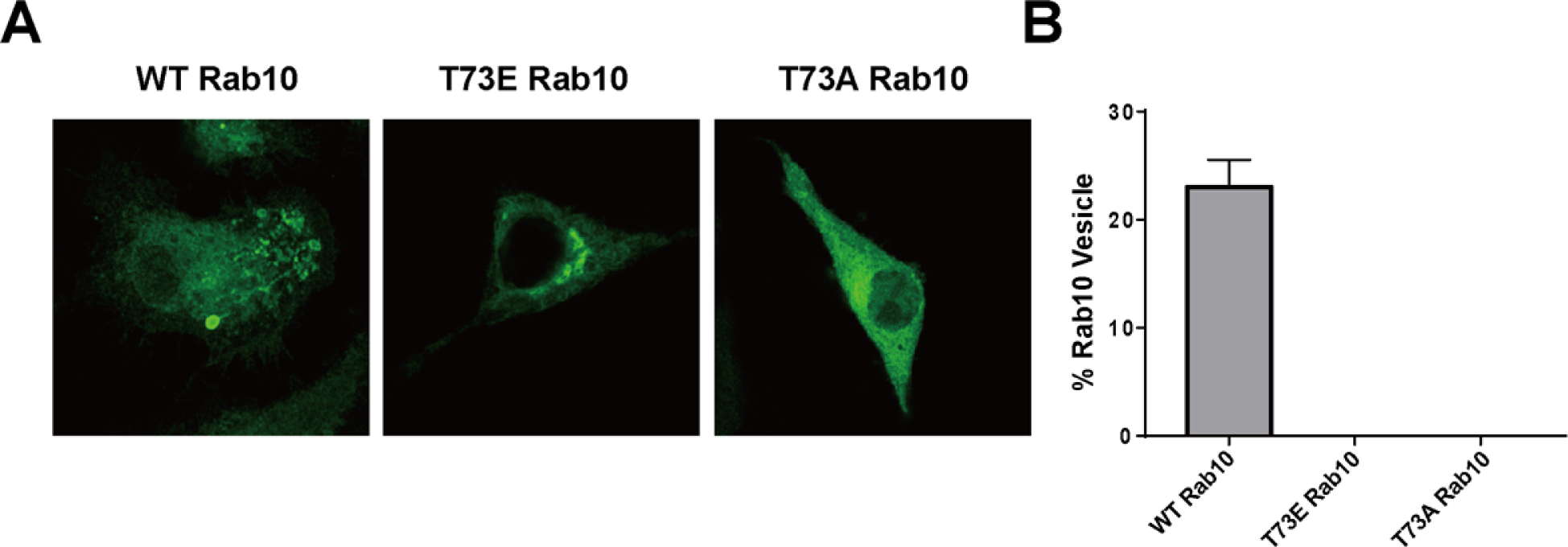
T73E Rab10 and T73A Rab10 do not show vesicular localization. D) Raw264.7 cells were transfected with eGFP(N-term)-WT or T73E or T73A mutant Rab10(show as green signal). Representative photomicrographs (from >20 images analyzed for each conditions from n=3 biologically independent experiments) are shown. Scale bars indicate 20 μm.

**Supplemental Figure 7.**
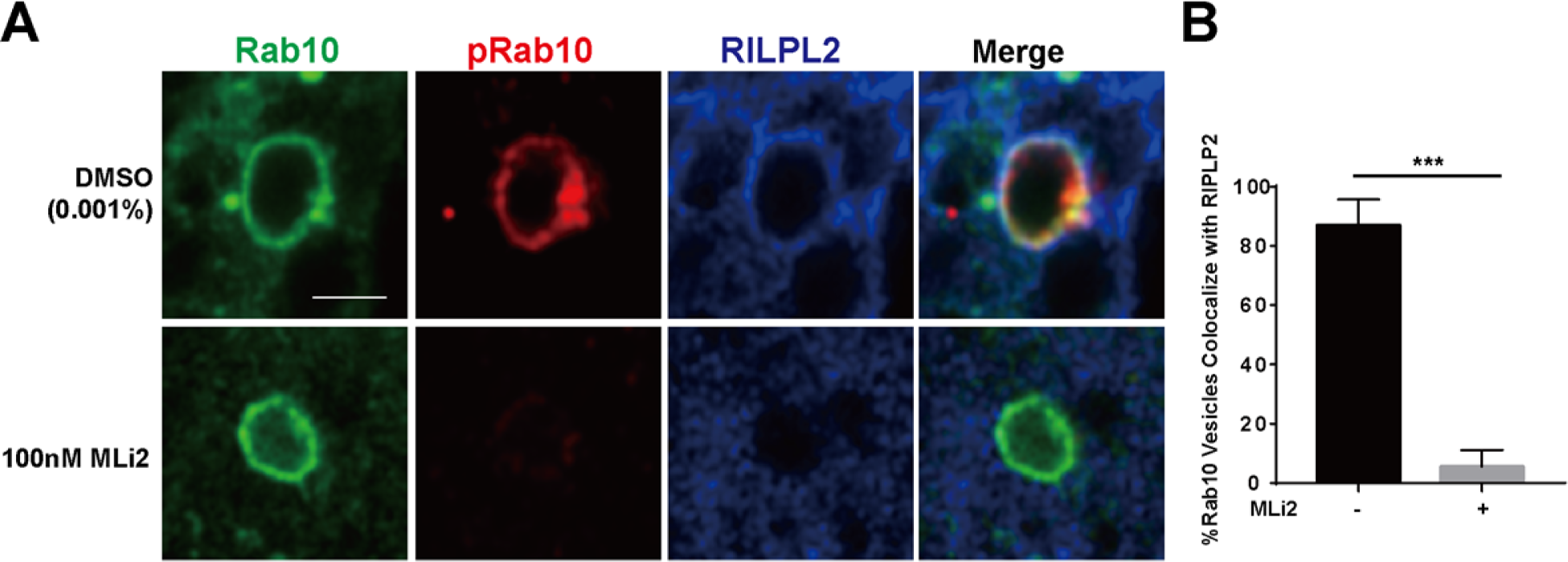
LRRK2 kinase activity is required for RILPL2-binding to Rab10 vesicles. **a,** Raw 264.7 cells were co-transfected with eGFP(N-term)-Rab10 and Flag(C-term)-RILPL2. Cells were immuno-stained with anti-Flag antibody for RILPL2 (shown in blue), and anti-pThr73 for pT73-Rab10 (shown in red). Representative photomicrographs (from >30 images analyzed for each condition across n=3 biologically independent experiments) are shown. Scale bar represents 1 μm. **b,** Calculated percentage of GFP-Rab10 and Flag(C-term)-RILPL2 double-positive vesicles, with or without MLi2 treatment, were calculated from >30 images analyzed for each condition across n=3 biologically independent experiments. Error bars are shown as ± SEM and significance was assessed by one-way ANOVA with Tukey’s post hoc test *p < 0.01.

**Supplemental Figure 8.**
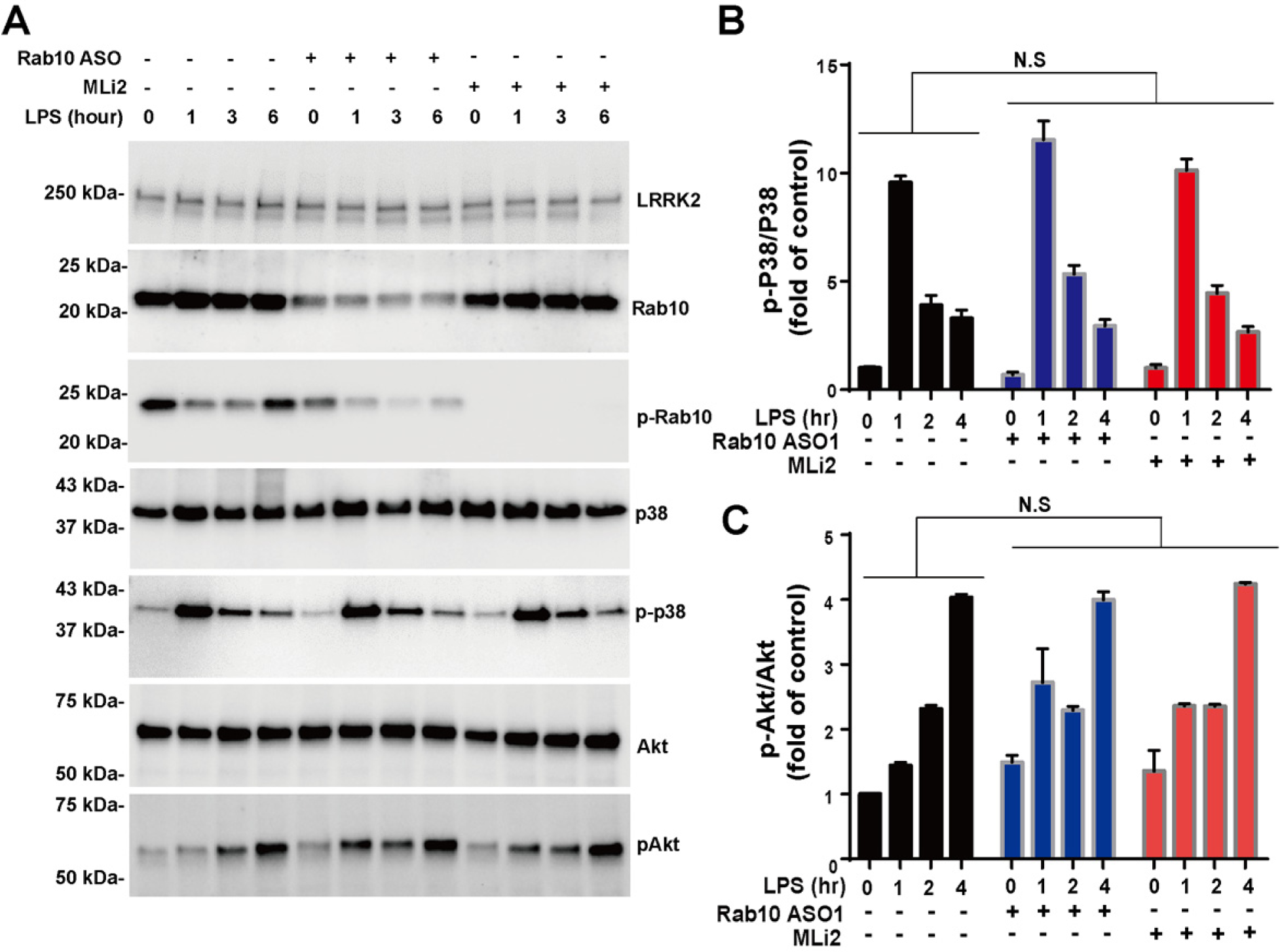
Rab10 knockdown has no effect on TLR4-stimulated MAPK activation. **a,** Primary mouse bone marrow-derived macrophage cells (mBMDM) from adult male WT-mLRRK2 BAC mice were treated with DMSO, 1 μM Rab10 ASO for 4 days, or 100 nM MLi2 for 12 hours before stimulated with 100 ng mL^-1^ LPS for the indicated time points. Representative immune-blots from n=3 biologically independent experiments are shown. **b,** Relative Phospho-p38 MAPK (Thr180/Tyr182) levels were normalized to total p38 and calculated as the fold increase compared to control naïve (no LPS) cells. **c,** Relative phospho-AKT (Ser473) levels were normalized to total AKT and calculated as the fold increase compared to control naïve (no LPS) cells. Columns indicate mean values, with error bars showing ± SEM; significance for each group (0, 1, 2, or 4 hours post-LPS) was assessed by a two-way ANOVA, with n.s. indicating p>0.5.

